# Selection for targeted therapy resistance leads to an indirect selection for higher phenotypic plasticity and enhanced evolvability to orthogonal stressors

**DOI:** 10.64898/2026.01.13.699073

**Authors:** Alicia Bjornberg, Aobuli Xieraili, Matthew Froid, Rowan Barker-Clarke, Jeff Maltas, Robert Vander Velde, Jamir Riffas, Berkley Gryder, Alexander RA Anderson, David Bassanta, Jacob Scott, Virginia A Turati, Andriy Marusyk

## Abstract

Acquired resistance to targeted therapies is a primary barrier to durable cancer control. Resistance frequently coincides with stem-like features and EMT-associated cell-state changes (partial EMT), yet whether these programs directly cause resistance or instead enable escape by increasing phenotypic plasticity remains debated. Integrating computational modeling, functional experimental assays, and lineage tracing, we investigated how cell-state plasticity shapes acquired resistance to ALK inhibition in ALK+ lung cancer models. Our results support a model in which drug exposure selects for cells with higher phenotypic plasticity, progressively increasing their representation as resistance emerges. Consequently, resistant populations show enhanced capacity to adapt to orthogonal therapeutic and environmental stressors and exhibit heightened metastatic potential. Bulk ATAC-seq showed that highly plastic cells have increased chromatin accessibility at regulators of EMT and stemness. Consistent with this, boosting plasticity via Yamanaka-factor induction or EMT-factor expression reduced ALKi sensitivity over time. In contrast, constraining plasticity (SOX2 knockdown or epigenetic inhibition) reduced long-term resistance outgrowth and prolonged ALKi response. Together, our results indicate that targeted therapy indirectly selects for cells with increased phenotypic plasticity, providing the substrate from which multifactorial resistance and metastatic competence evolve. Further, it suggests that constraining plasticity could delay resistance and extend response durability.

## INTRODUCTION

Targeted therapies, particularly small-molecule inhibitors designed to exploit oncogenic signaling dependencies, have transformed the treatment landscape for advanced cancers^1–3^. In subsets of patients, these agents induce profound and durable initial responses. However, in most cases, these agents fail to eradicate disease, and residual cells eventually acquire resistance, leading to relapse^4^. Defining the principles that govern resistance emergence is therefore essential for extending response durability.

Mechanistically, acquired resistance has traditionally been framed as a mutational problem: on-target mutations that limit the drug binding or off-target mutations that activate bypass signaling restore proliferation in the presence of drug^5^. Yet many tumors relapse without an immediately identifiable dominant resistance driver, suggesting that resistance can involve non-genetic, cell-state adaptations in addition to canonical genetic routes^6,7^. Across systems, therapy exposure is accompanied by reversible transcriptional and chromatin remodeling, often coupled to stress-response programs, creating a permissive backdrop from which more stable resistant phenotypes can emerge over time^8–10^.

In our prior work with experimental models of ALK-positive non-small lung cancers (ALK+ NSCLC), we found that resistance accrued through a multifactorial combination of genetic and non-genetic adaptations within the same cells, following inhibitor-specific trajectories in adaptive and epigenetic landscapes rather than a single dominant resistance event ^11^. Similar multi-step resistance trajectories have been reported in independent systems ^12,13^.

These state-based models intersect with broader programs linked to plasticity, including EMT- and stemness-associated transcriptional networks (henceforth EMT/stemness), which have been repeatedly observed in resistant tumors and metastasis^14,15^. EMT/ stemness signatures are often interpreted as proximal resistance mechanisms^16–18^; however, we previously showed that experimentally activating EMT is insufficient to confer complete resistance^12^, motivating the alternative possibility that these programs mark increased capacity for adaptive cell-state transitions, particularly in settings where resistance emerges gradually through multiple adaptations.

To dissect the relationship between plasticity and adaptive capacity, we combined *in silico* modeling with molecular and functional analyses of therapy-naïve and therapy-resistant NSCLC cells. We show that acquired ALK inhibitor (ALKi) resistance is accompanied by broader ability to adapt to orthogonal pharmacological and environmental stressors and by increased metastatic potential.

Our data supports a model in which this broader adaptability stems from indirect selection for increased phenotypic plasticity under therapeutic pressure. We further show that this effect is most pronounced when resistance evolves through a multifactorial, stepwise trajectory. In line with this, adaptive potential can be increased or reduced by genetic and pharmacological perturbations. Together, these results suggest that limiting phenotypic plasticity may prolong response durability by delaying resistance emergence.

## RESULTS

### Independent ALKi resistance trajectories converge on EMT/stemness and cross-stressor adaptability

Our previous work in an experimental model of ALK+ NSCLC showed that the development of resistance to clinically relevant concentrations of multiple ALKi is characterized by the gradual accumulation of resistance-conferring molecular changes. Whereas mutational changes contributed to this process, adaptation was dominated by expression-level, epigenetic changes. Surprisingly, even though resistance to any given ALKi was associated with cross-resistance to different ALKi, evolution of resistance to distinct ALK inhibitors was characterized by predictively distinct, inhibitor-specific evolutionary trajectories, and mediated by distinct combinations of resistance-promoting mechanisms^11^. Building on these experiments, we asked, despite these differences, whether there is a common denominator to ALKi resistance. To this end, we focused on resistance to the two most clinically relevant ALKi, alectinib and lorlatinib. In our experiments, resistant variants of the H3122 cell line were derived by continuous exposure to clinically relevant drug concentrations until stable resistance emerged, yielding lorlatinib-evolved (erLor) or alectinib-evolved (erAlec) resistant cells (collectively erALKi) (**Fig. 1A**).

**Figure 1.**
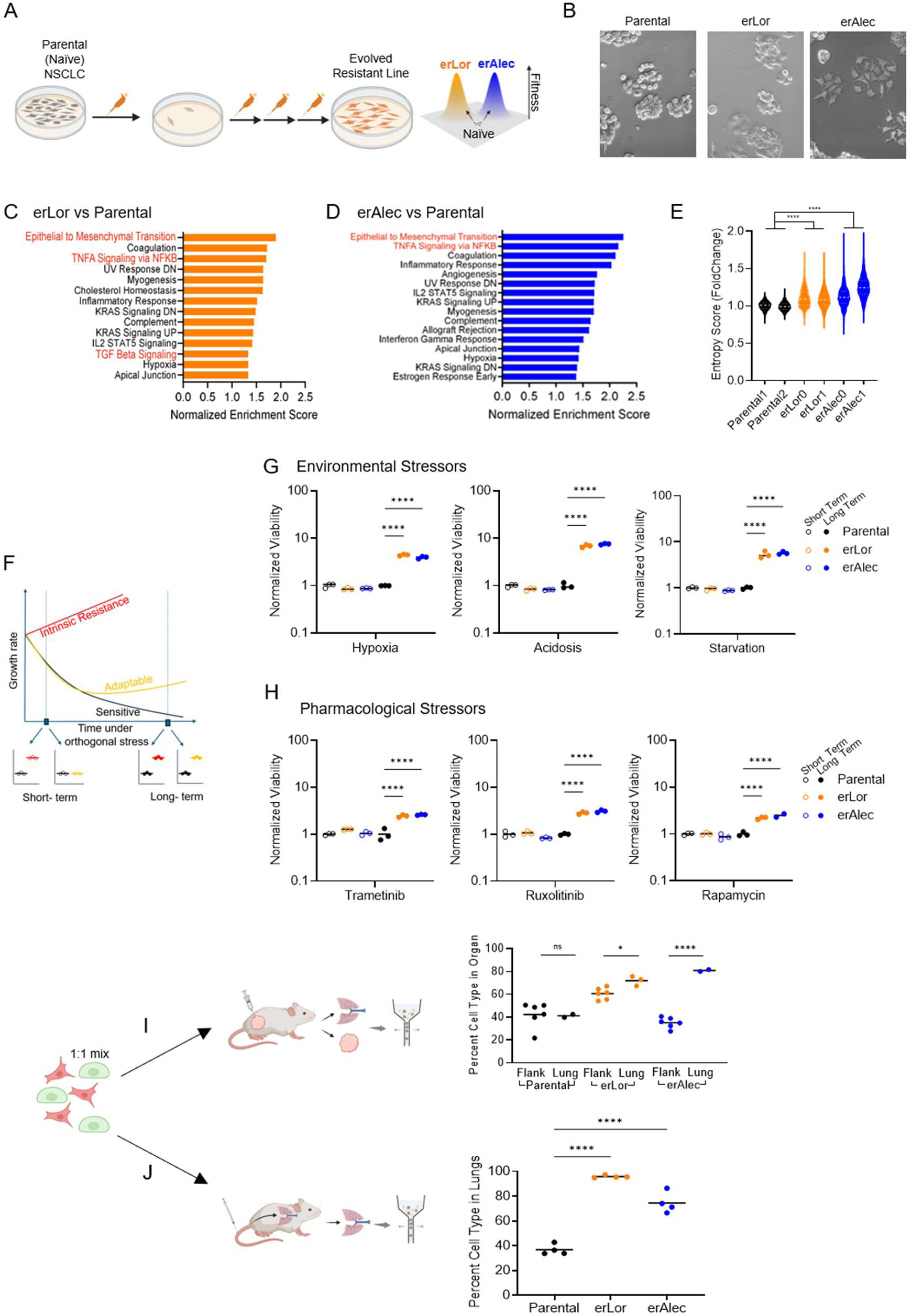
Independent treatment of NSCLC with different TKIs demonstrates convergence towards similar resistance phenotypes. **A.** Schematic of the experimental design to derive evolved resistant cells to lorlatinib (erLor) and alectinib (erAlec). **B.** Representative bright-field images of Parental, erLor, and erAlec cells in normal media at 20x magnification (Scale bar 50μm). **C.** RNA-sequencing analysis of significantly enriched pathways from Gene Set Enrichment Analysis (GSEA) of hallmark gene sets in erLor versus Parental cells or (**D**) erAlec vs Parental cells. Pathways with ≥1.33 normalized enrichment score (NES) shown. **E.** Violin plot of Gini score of transcriptional entropy for RNA sequencing data of two biological replicates each of Parental, erLor, and erAlec. One-way ANOVA with Tukey’s multiple comparisons test. **F.** Schematic of expected results of short and long term experiments contrasting intrinsic and acquired resistance. **G.** Quantification of viability staining after short term or long term treatment with environmental conditions hypoxia (200nM CoCl2), acidosis (pH of 5.5), or starvation (1:10 dilution) normalized to growth under normal media. Mean and three replicates shown. Ordinary One-way ANOVA with Tukey’s multiple comparisons test was used; only comparisons to the parental sample within each time period are shown. **H.** Quantification of viability staining after short term or long term treatment with trametinib (15nM), ruxolitinib (20μM), or rapamycin (10μM) normalized to vehicle control (DMSO, 0.025%). Mean and three replicates shown. Ordinary One-way ANOVA with Tukey’s multiple comparisons test was used; only comparisons to the parental sample within each time period are shown. **I.** Left: schematic of experimental design where mixed populations of Parental GFP cells and mCherry, either erLor or erAlec, cells were injected subcutaneously. Both the primary subcutaneous site and lung metastatic site were collected and cells enumerated via FACS. Right: percent of each cell type found in given organ based on FACS analysis after 27 days post injection. Unpaired t-tests shown. **J.** Left: schematic of experimental design where a mixed populations of Parental GFP+ cells and mCherry+, either erLor or erAlec, cells were injected via tail vein. Right: percent of each cell line found in the lungs after 27 days post injection. One-way ANOVA with Tukey’s multiple comparisons test was used; only comparisons to Parental shown.

Notably, erALKi cells displayed a more mesenchymal morphology compared to their treatment-naïve counterparts (**Fig. 1B**). Further, transcriptomic profiling and gene set enrichment analysis (GSEA) showed that erAlec and erLor cells shared activation of TNF-α and TGF-β signaling (**Fig. 1C–D, Fig. S1A–B**), pathways implicated in EMT, stemness, and metastasis^19–25^. While EMT is often considered to be a direct resistance mechanism^16–18^, our previous studies demonstrated that EMT induction, per se, have modest to negligible effect on drug sensitivity in common cell viability assays^12^. Given the association of intermediate EMT with higher phenotypic plasticity and stemness, we hypothesized that the convergent EMT-enrichment in erALKi cells might reflect selection for cells with higher capability of undergoing adaptive cell-state transitions under sustained therapeutic selection pressures.

To test this hypothesis, we first asked whether erALKi cells display higher transcriptional entropy, which can be captured with the Gini index, a metric previously used to quantitatively infer differentiation potential and plasticity in normal and cancer stem cells^26,27^. Indeed, erALKi cells exhibited significantly higher entropy compared with treatment-naïve cells, consistent with the elevated EMT/stemness (**Fig. 1E**). Next, we sought to evaluate our hypothesis more directly in a functional assay. We reasoned that if EMT reflects a higher plasticity cell state, the erALKi cells should be more capable of adapting to orthogonal stressors, without necessarily showing higher fitness in short term assays. In this case, adaptability can be distinguished from intrinsic resistance by contrasting the initial sensitivity of therapy-naïve and erALKi cells to orthogonal stressors, with their sensitivity after prolonged incubation under stress once cells have had an opportunity to adapt. In this experimental design, higher plasticity-mediated adaptive potential could be identified as a higher viability in long-term, but not short-term assays (**Fig. 1F**).

We started by contrasting short- and long-term sensitivity of erALKi and naïve H3122 cells to microenvironmental stresses commonly encountered within tumors: hypoxia, acidosis, and nutrient starvation^28,29^. After short-term exposure (3 days), resistant and naïve cells showed similar survival and growth, indicating that EMT does not confer an intrinsic resistance to these stressors. However, the prolonged exposure (≥10 days; specific duration chosen based on the stressor-specific adaptation timelines), erALKi cells displayed markedly higher viability (**Fig. 1G**), consistent with a higher adaptive potential. Next, we asked whether erALKi cells are more capable of adapting to clinically relevant orthogonal signaling inhibitors targeting MEK1/2 (trametinib), JAK1/2 (ruxolitinib), and mTOR (rapamycin)^30–33^. Similar to the observations with environmental stressors, while early responses were comparable, resistant populations progressively adapted and displayed increased growth relative to naïve cells (**Fig. 1H, Fig. S1C**). These data show that EMT phenotypes of erALKi cells are associated with a generalized, time-dependent capacity to adapt to unrelated pharmacological and environmental stressors, rather than an intrinsic resistance to orthogonal stressors.

We next asked whether this adaptive potential extends to an enhanced ability to adapt to new tissue environments. Metastatic colonization represents a stringent selective bottleneck that requires plasticity-mediated adaptation^34^. To test whether ALKi resistant states exhibit broader adaptive capacity, we injected 1:1 mixtures of mCherry-labeled resistant cells (erLor or erAlec) and GFP-labeled naïve cells subcutaneously into the flanks of NSG mice. While the primary tumors maintained the initial 1:1 ratio, consistent with the lack of engraftment and growth differences between naïve and erALKi cells, analysis of spontaneous lung metastases by fluorescence-activated cell sorting (FACS) revealed enrichment of resistant cells, with two-fold more mCherry+ (erLor and erAlec) than GFP+ (naïve) cells recovered from metastatic lesions (**Fig. 1I**). Control mice injected with naïve cells alone and comparable flank engraftment excluded fluorophore bias and differences at the primary site, indicating that enrichment was specific to the metastatic setting. To distinguish between enhanced dissemination from the primary tumor versus superior adaptability at the metastatic site, we bypassed local invasion and intravasation by injecting resistant and naïve cells directly into the circulation via the tail vein^35^. Resistant cells again outcompeted naïve cells by two- to three-fold in lung colonization (**Fig. 1J, Fig. S1D**), demonstrating that resistant cells adapt more effectively to the lung microenvironment, a property consistent with enhanced plasticity.

Together, these data show that ALK inhibitor resistance converges on EMT/stemness-associated programs, characterized by increased transcriptional entropy and a higher capacity for adaptation towards orthogonal stressors and new tissue contexts.

### Multi-step resistance dynamics predict progressive enrichment of rare high-plasticity cells

The convergence of ALKi resistance on phenotypes characterized by elevated phenotypic plasticity suggests that selection for ALKi resistance leads to indirect selection for enhanced phenotypic plasticity. We speculated that this strong convergence might reflect the complexity and multifactorial nature of resistance dynamics revealed in our recent studies^11,12^, where a multitude of steps towards resistance might strongly amplify the indirect selection for higher adaptability. To test this prediction, we developed an *in silico* model of heterogeneous tumor populations contrasting the selective advantage of increased adaptability under single-hit and multi-step resistance acquisition scenarios (**Fig. 2A**).

**Figure 2.**
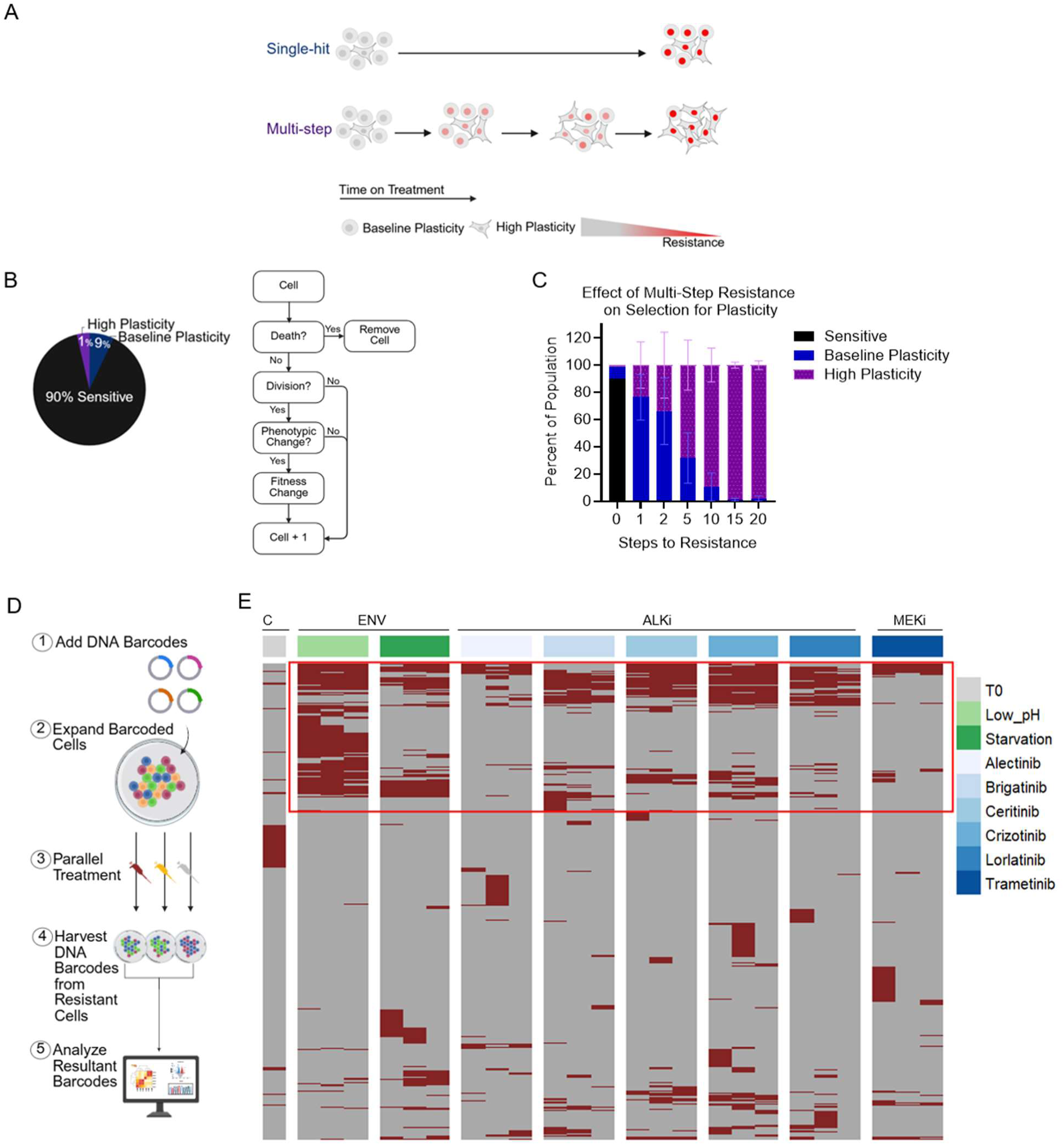
Treatment and exposure to harsh environments drive gradual selection for rare cells with higher adaptability. **A.** Schematic of hypotheses. **B.** Left: schematic of initial simulation conditions. Right: schematic of agent-based model (ABM) algorithm. **C**. Cell fraction of each population after 600 timesteps based on the number of steps to resistance modeled in each simulation. Mean ± SD shown of 10 simulations. **D.** Schematic of experimental design for barcode libraries. **E.** Unsupervised hierarchical clustering of barcodes from CloneTracer library in each sample based on presence in given treatment. Each row represents a barcode. Each column represents a replicate of a condition. n= 1 for T0, n=3 for all other arms. C = control, T0= time zero, initial time point; ENV = environmental stressor, ALKi= ALK inhibitor, MEKi= MEK inhibitor. DMSO (0.1%), alectinib (0.5μM) lorlatinib (0.5μM), crizotinib (0.5μM), ceritinib (0.1μM), brigatinib (0.1μM), trametinib (0.1μM), Low_pH pH of 6, starvation 1:5 media:PBS.

To capture both spatial and well-mixed dynamics, we used two approaches: a spatial agent-based model (ABM) (**Fig. S2A**) and a non-spatial ordinary differential equation (ODE) model (**Fig. S2B–C**, **Supplementary Methods**). In both ABM and ODE implementations, cells differed in an abstract plasticity parameter that sets the probability of developing phenotypic changes that increase fitness under treatment. Cells could transition towards fitter resistant states stochastically, allowing us to track subpopulation dynamics under treatment over time.

In our simulations we sought to recapitulate a scenario of therapeutic responses of a therapeutically sensitive neoplastic population composed of cells with varying adaptation potential. Of the 10,000 cells initialized in the simulation, the majority (90%) are incapable of adaptation (phenotypic-change rate = 0); these are meant to reflect cells that are effectively eliminated by therapy. The remaining 10% are capable of acquiring resistance-promoting adaptations: 9% exhibit baseline adaptability (1x change rate), while just 1% exhibit higher adaptability (2x change rate) (**Fig. 2B**). The relatively modest difference in phenotypic change rates was chosen deliberately; as larger differences should inevitably amplify the expected effects. At each step, cells could die, divide, or undergo a phenotypic change representing cumulative genetic and/or epigenetic modifications without specifying molecular origin (**Fig. 2B**). Key parameters were systematically varied, including the initial frequencies of plastic subpopulations and the nature of treatment applied. We modeled cytostatic therapy by reducing proliferation (**Fig. S2B**) and cytotoxic therapy by increasing death (**Fig. S2C**).

Whereas partial selection for cells with higher plasticity potential was observed under the single-hit scenario, these resistant populations were dominated by the descendants of the initially more numerous cells with baseline plasticity. In contrast, across parameter regimes, the multi-step framework progressively enriched subpopulations with elevated plasticity (**Fig. 2C, Fig. S2A–D**). Together, our *in silico* simulations indicate that i) evolution of resistance is associated with an indirect selection for cells with higher adaptability potential and ii) that this selection is strongly augmented under a scenario of multi-step resistance.

### Individual NSCLC clones can be classified into adaptation potentials with distinct functional and molecular profiles

Next, we sought to examine the existence of empirical evidence for the simulation’s key assumption, that treatment-naïve populations contain rare lineages with elevated adaptability. To this end, we performed high-complexity lineage tracing using two independent DNA barcode libraries (CloneTracer^36^ and CloneSweeper^37^). Each barcoded population was challenged with a panel of kinase-pathway inhibitors and microenvironmental stressors (**Fig. 2D**). Barcode composition before and after each challenge was compared, and enrichment relative to baseline composition was used to identify lineages that expanded across conditions.

Across both libraries, we observed a rare subset of barcodes that expanded under multiple conditions, with partial reproducibility across replicates, consistent with stochastic selection and context-dependent adaptive solutions rather than a single fixed “winner” lineage (**Fig. 2E, Fig. S3A–D**). Specifically, we observed reproducible structure in which stressors tend to co-select overlapping sets of lineages. For example, in one dataset, barcodes enriched under ceritinib, crizotinib, lorlatinib, and brigatinib clustered separately from those enriched under alectinib and trametinib, or low pH and nutrient starvation (**Fig. S3B**). In the other library, barcodes selected by alectinib and lorlatinib were grouped apart from those expanded under acidic conditions (**Fig. S3D**). While our lineage tracing experiments indicate a complex scenario where distinct subpopulations vary in their ability to adapt to distinct stressors, they also support the notion of existence of a subpopulation of cells with an enhanced general adaptability within therapy-naïve H3122 cells.

To test for this inference more directly, we derived single-cell clones from treatment-naïve H3122 cells (**Fig. 3A**) and contrasted their ALKi adaptation potential. Using flow sorting, we seeded single cells into individual wells of 96-well plates and verified single-cell seeding by microscopy. From 127 sorted cells, 21 clonal colonies were established and then challenged independently with clinically relevant concentrations of ALK inhibitors (alectinib, lorlatinib, crizotinib, ceritinib, brigatinib). Clonal responses were quantified using long-term clonogenic outgrowth assays after sustained drug exposure, with resistant colonies visualized by crystal violet staining (**Fig. 3B-C**). Whereas baseline proliferation and short-term ALKi sensitivity showed no substantial variability, long-term clonogenic outgrowth varied markedly across clones, revealing clear differences in the ability to adapt under continued ALK inhibition (**Fig. 3D, Fig. S4A-B**). Two clones capable of clonogenic outgrowth across all ALKi were denoted high adaptation-potential (HAP). Eleven clones that failed to establish colonies under any ALKi were denoted low adaptation-potential (LAP). The remaining eight clones displayed intermediate behavior, with clonogenic outgrowth under some but not all ALKi, and were denoted intermediate adaptation-potential (IAP). This distribution is consistent with our barcoding data and prior reports indicating that resistance originates from a rare fraction of the therapy-naive tumor cell populations^6,11,12,36^.

**Figure 3.**
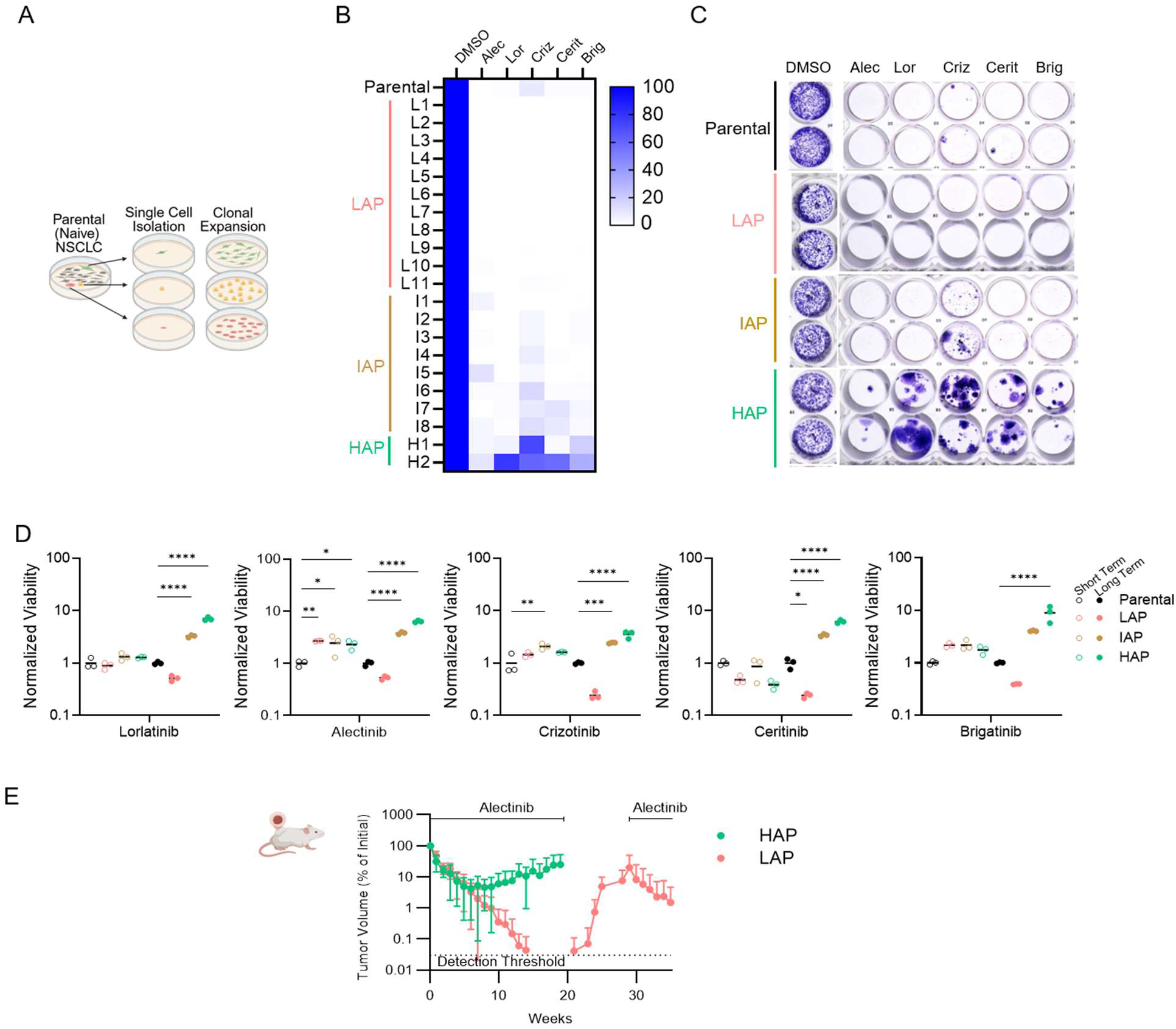
Single-cell-derived clones can be classified into adaptation potentials with distinct functional and molecular profiles. **A.** Schematic of experimental categorization of single-cell derived clones. **B.** Heatmap representing Image J crystal violet quantification as a percent confluence of each cell line compared to vehicle treatment (DMSO) for up to three months in treatment. Drug treatment displayed as rows, clone number listed as columns for the 21 clones that expanded from single-cell sorting, with final classifications as low, intermediate, or high adaptation-potential. **C.** Crystal violet staining as quantified in Fig. 3B of Parental, HAP, IAP, and LAP cells in DMSO 0.05%, alectinib 0.5μM, lorlatinib 0.5μM, crizotinib 0.5μM, ceritinib 0.1μM, or brigatinib 0.1μM. **D.** Cell Titer Glo quantification of representative clones from each category of LAP, IAP, or HAP. Percent viability after 3 days (short term) or 21 days (long term) normalized to growth under vehicle control. Mean and three replicates shown. Ordinary One-way ANOVA with Tukey’s multiple comparisons test was used; only comparisons to the parental sample within each time period are shown. **E.** Volumetric traces of HAP and LAP H3122 xenograft tumors on treatment with 25mg/kg alectinib for 20 weeks, followed by a treatment break for LAP tumors for 9 weeks, followed by re-treatment for LAP tumor-bearing mice for 5 weeks. Mean ± SD shown. P values using mixed effect model in Supplementary Table 1 (n= 8 tumors per group).

To test whether the adaptation phenotypes were stable over time in culture, naïve LAP, HAP, and parental cells were passaged and plated weekly into clonogenic assays (**Fig. S4C**).

Cells were then treated with ALKi for four weeks to allow resistant colony formation, followed by crystal violet staining and quantification. The phenotypes remained stable over eight weeks of passaging: HAP clones consistently formed larger and more numerous resistant colonies, whereas LAP clones failed to generate colonies under ALKi (**Fig. S4D**).

The parental population formed colonies at intermediate frequency, as expected from a heterogeneous mix of lineages. These results demonstrate that the observed differences in adaptation potential are stable over extended culture and are not altered by routine serial passaging.

Next we asked whether the variability in ALKi adaptation potential extends towards *in vivo* responses. To this end, xenograft tumors were generated from LAP and HAP cells and subcutaneously implanted into NSG mice. Consistent with the *in vitro* assays, we did not observe differences in the initial alectinib responses between LAP and HAP xenografts.

However, around 8 weeks of treatment responses started to diverge, with HAP tumors developing resistance while LAP tumors continued to regress (**Fig. 3E**). To assess whether LAP tumors were fully eliminated, the treatment was discontinued at week 21. Treatment discontinuation triggered tumor relapse, indicating the survival of viable tumor cells within microscopic residual disease. After relapsing tumors reached pre-treatment sizes, alectinib treatment was resumed. Notably, relapsed tumors displayed strong responses, similar to the responses of treatment-naïve LAP tumors, indicating that LAP tumor cells failed to develop measurable resistance (**Fig. 3E**).

We then asked whether the enhanced ALKi adaptive potential of HAP cells reflected an enhanced adaptability in general. To this end, we used our functional assay pipeline and panel of orthogonal stressors initially developed to uncover the enhanced adaptability potential of ALKi cells (**Fig. 1F-H**). We found that while the initial sensitivity of HAP cells to orthogonal stressors was similar to the sensitivity of LAP, IAP and parental cells, the HAP cells were more capable of developing resistance over time (**Fig. 4A-B**). Finally, we asked whether, similar to ALKi cells, HAP cells are more capable of metastatic colonization. To this end, LAP, HAP, and parental cells were engineered to express Cypridina luciferase^38^, enabling longitudinal vital monitoring of metastatic burden after tail-vein injection (**Fig. 4C**). Within two weeks, the luciferase signal dropped below the detectability threshold in all of the injected mice, consistent with the expectation of substantial selection bottleneck. By 15 weeks, all mice injected with HAP cells exhibited strong luciferase signal, indicating outgrowth of metastatic tumors, whereas plasma luciferase levels in LAP-injected mice remained below the detection threshold (**Fig. 4D**). Parental cells displayed an intermediate phenotype: whereas 3/4 injected animals had detectable luciferase signal, the kinetics were delayed compared to HAP cells (**Fig. 4D**), consistent with their heterogenous composition containing only a minority of high adaptation-potential clones. Examination of tissues at point of euthanasia validated presence of macroscopic lung and liver tumors in the animals with detectable luciferase signals (**Fig. S6A**), while no detectable tumors were observed in animals seeded with LAP cells. Together, these results support the notion that therapy-naïve tumor cells contain subpopulations with an increased general adaptation potential.

**Figure 4.**
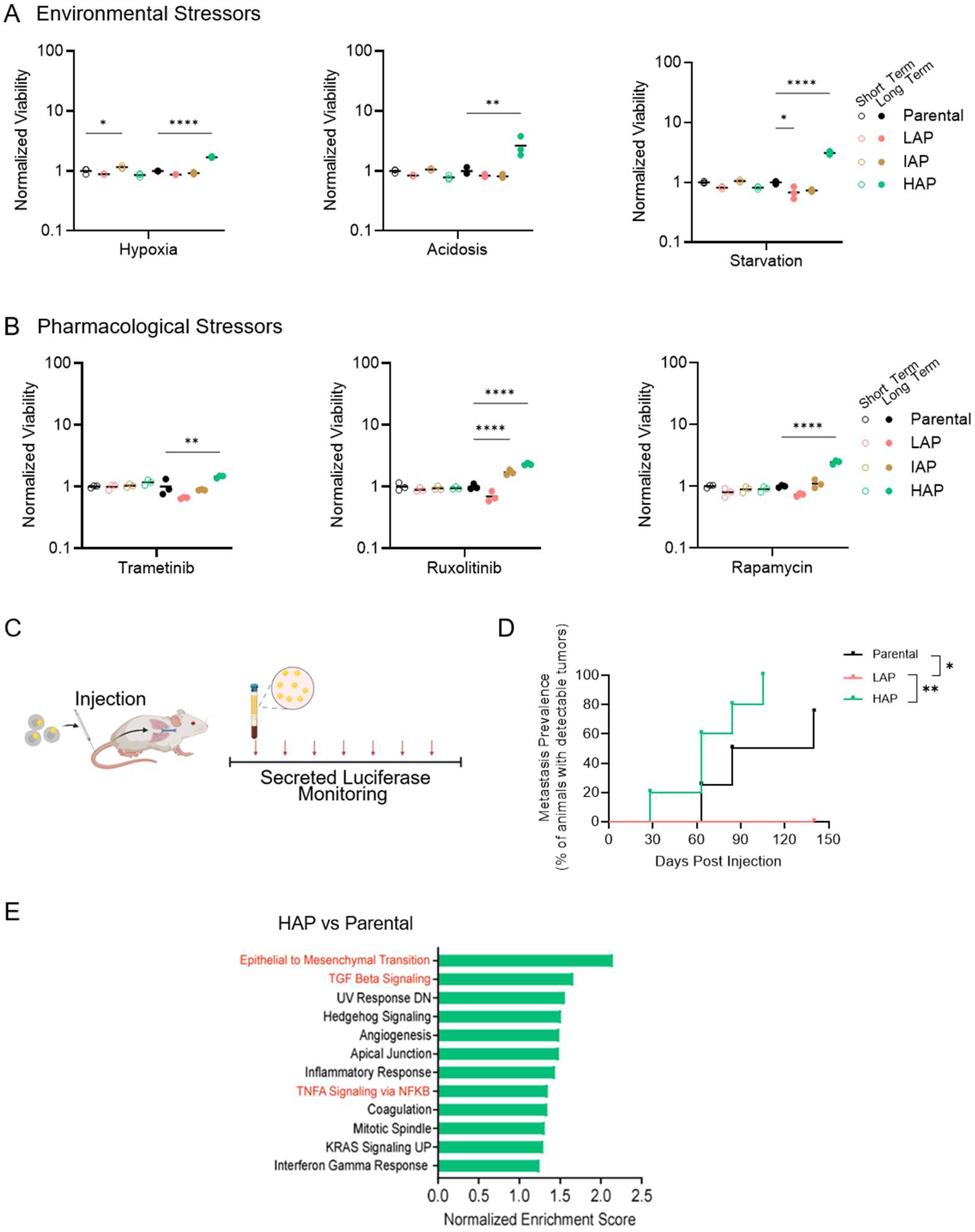
TKI adaptation-potential extends to metastatic potential and multi-stress resistance. **A.** *In vitro* quantification of crystal violet viability staining of representative cell lines of Parental, LAP, IAP, or HAP in hypoxia (200nM CoCl_2_), acidosis (5.5 pH), or starvation (1:5 media:PBS) normalized to growth under normal media after 3 days (short term) or 16 days (long term). Mean and three replicates (short term) or two replicates (long term) shown. Ordinary One-way ANOVA with Tukey’s multiple comparisons test was used; only comparisons to the parental sample within each time period are shown. **B.** *In vitro* quantification of crystal violet viability staining of representative cell lines of Parental, LAP, IAP, HAP, erLor, or erAlec in trametinib (15nM), ruxolitinib (20μM), rapamycin (10μM) normalized to growth under vehicle control (DMSO, 0.025%) after 3 days (short term) or 22 days (long term). Mean and three replicates shown. Ordinary One-way ANOVA with Tukey’s multiple comparisons test was used; only comparisons to the parental sample within each time period are shown. **C.** Schematic of metastasis experiment with continuous monitoring via luciferase assay of blood plasma. **D.** Cumulative Incidence plot of percent of animals in each group that developed metastasis, as measured by luciferase activity in plasma. n=5 for LAP and HAP groups, n=4 for Parental group. Logrank (Mantel-Cox) test shown. **E.** Significantly enriched pathways from Gene Set Enrichment Analysis (GSEA) of hallmark gene sets in HAP versus Parental cells from RNA sequencing. Pathways with ≥1.25 normalized enrichment score (NES) shown.

### Chromatin accessibility informs therapeutic vulnerabilities to modulate adaptation potential

To identify the mechanistic underpinning of differences in the adaptive potential, we started with characterization of transcriptional programs. To this end, we performed bulk RNA-seq on LAP, IAP, and HAP clones. Unsupervised clustering and PCA grouped clones by class (**Fig. S5A–B**), indicating that LAP/IAP/HAP subpopulations correspond to stable, molecularly distinct transcriptional states. LAP clones showed the greatest inter-class heterogeneity, whereas HAP clones displayed more consistent profiles. GSEA revealed that, similarly to cells with evolved resistance, HAP clones upregulated EMT- and stemness-associated pathways, including TGF-β, Hedgehog, and TNFα/NF-κB (**Fig. 4E, Fig. S5C**)^19–25,39^. Consistent with these profiles, LAP clones retained the rounded cobblestone morphology of parental cells, whereas HAP clones displayed a more mesenchymal phenotype (**Fig. S5D**).

Chromatin accessibility can shape adaptation by regulating transcriptional responses to environmental change^40^. To test whether adaptation potential is associated with a distinct regulatory landscape we next profiled chromatin accessibility by bulk ATAC-seq^41^. HAP cells displayed increased accessibility at developmental, cell-identity regulators, and enrichment of EMT/stemness-associated motifs, including KLF4 and SOX family members (**Fig. 5A–B**, **Fig. S6B-C**). Whereas the reading depth of our bulk RNA seq analyses was insufficient to detect the expression of these transcriptional factors, the ATAC-seq analyses support the notion that the enhanced expression of EMT programs in HAP clones is associated with higher stemness.

**Figure 5.**
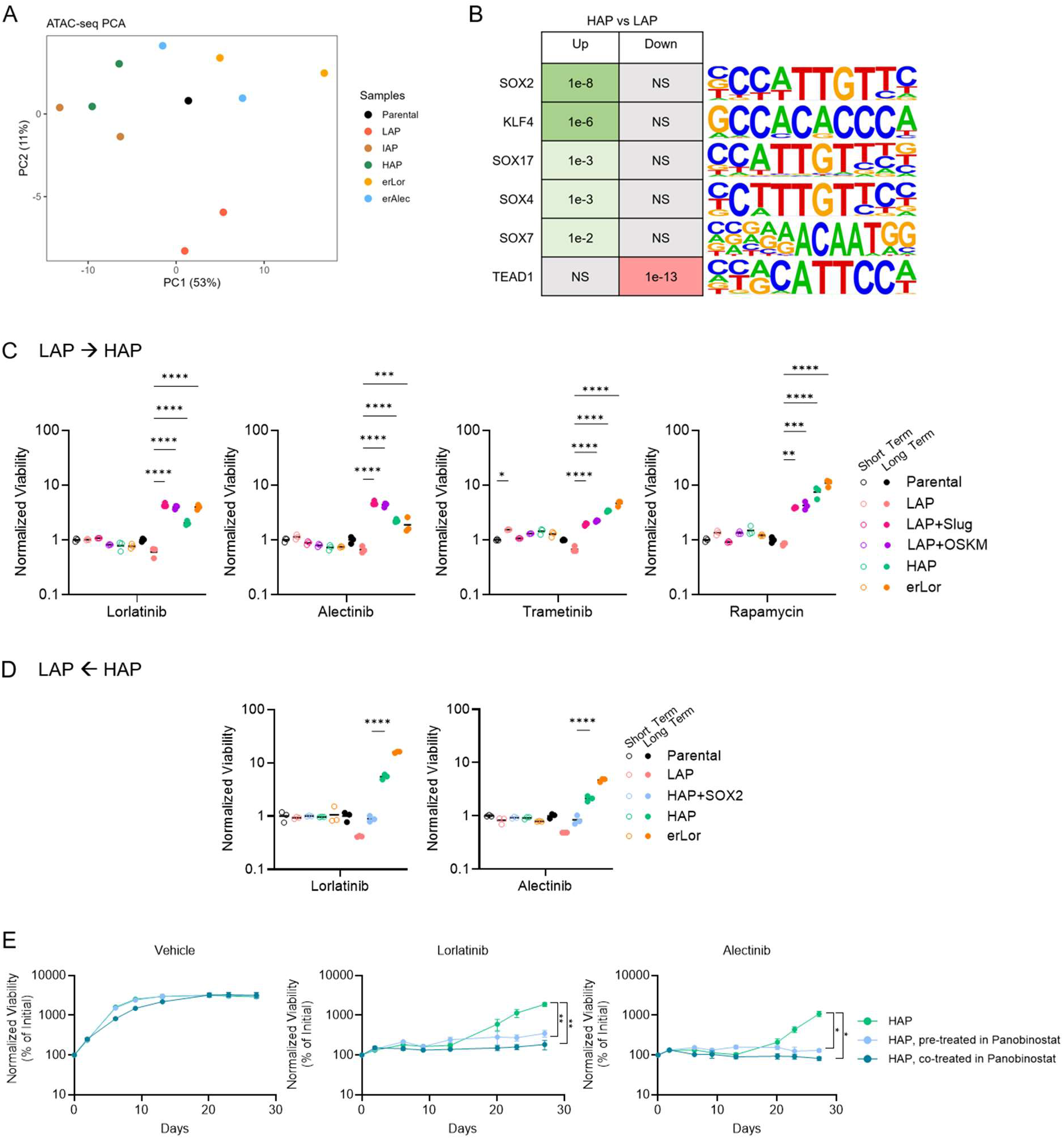
Adaptation-potential can be manipulated. **A.** PCA of top 500 variable genes from bulk ATAC- sequencing. **B.** HOMER known motif enrichment analysis of motifs enriched in HAP cells compared to LAP cells in relevant transcription factors. P -value shown in table. NS, not statistically significant. **C.** Quantification of crystal violet viability staining of Parental, LAP, HAP, LAP + Slug or LAP + OSKM cells in lorlatinib (0.5μM), alectinib (0.5μM), trametinib (15nM), or rapamycin (10μM) normalized to growth in vehicle control (DMSO, 0.025%), after 3 days (short term) or 4 weeks (long term). Mean and three replicates shown. Ordinary One-way ANOVA with Tukey’s multiple comparisons test was used; only comparisons to the LAP sample within each time period are shown. **D.** Quantification of crystal violet viability staining of Parental, LAP, HAP, HAP + shSOX2, and erLor cells in lorlatinib (0.5μM), or alectinib (0.5μM) normalized to vehicle control (DMSO, 0.025%) after 3 days (short term) or 4 weeks (long term). Mean and three replicates shown. Ordinary One-way ANOVA with Tukey’s multiple comparisons test was used; only comparisons between HAP and HAP+shSOX2 samples within each time period are shown. **E.** Quantification by live cell imaging of percentage of area covered by cells treated with vehicle (DMSO, 0.025%; left), lorlatinib (0.5μM; middle), or alectinib (0.5μM; right) with pre- treatment of co-treatment with Panobinostat (2nM). Two-way ANOVA with repeated measurements and Tukey’s multiple comparison in Supplementary Table 1; comparison to HAP shown.

We next tested whether adaptation potential can be shifted in either direction by perturbing cell-state regulators. To increase adaptation potential in LAP cells, we induced stemness via expression of Yamanaka factors (Oct4, SOX2, KLF4, c-MYC; OSKM)^42^ and, in parallel, induced EMT via overexpression of Slug (SNAI2)^12,43–46^. In both cases, these manipulations reduced sensitivity to ALKi over time in long-term assays (**Fig. 5C**) and were accompanied by a morphological shift toward a more mesenchymal phenotype (**Fig. S6D**), consistent with increased capacity to adapt under sustained drug challenge.

Conversely, we tested whether the high adaptation-potential phenotype could be dampened by limiting plasticity-associated regulators in HAP cells. Guided by our ATAC-seq results, which showed increased accessibility and motif enrichment for SOX family regulators in HAP cells, we performed shRNA-mediated knockdown of SOX2 (**Fig. S6E**). SOX2 knockdown did not restore acute ALKi sensitivity or alter short-term growth, but it significantly reduced the capacity to develop resistance in long-term assays (**Fig. 5D**).

Together, these bidirectional perturbations show that adaptation potential is experimentally modifiable, enhanced by enforcing stemness/EMT programs in LAP cells and reduced by limiting a plasticity-sustaining regulator in HAP cells, with corresponding effects on long-term resistance emergence (**Fig. 5C–D**).

Although direct pharmacological targeting of transcriptional factors is currently not feasible, because HAP and LAP cells show comparable short-term sensitivity to ALKi, we hypothesized that epigenetic inhibitors could limit adaptive phenotypic cell state transitions in HAP cells; thereby locking them in a drug-sensitive state. To test this, we treated HAP cells with either the histone deacetylase (HDAC) inhibitor panobinostat^47,48^ or the histone acetyltransferase (HAT) inhibitor A485^49^. Doses were selected to avoid direct cytotoxicity while producing a measurable stress response, as reflected by decreased NRF2 (*NFE2L2*) expression (**Fig. S6F**). Cells were then plated into ALKi either untreated, after 7-day pretreatment with an epigenetic inhibitor, or under continuous co-treatment with epigenetic inhibitor plus ALKi (**Fig. S6G**). In both pretreated and co-treated groups, panobinostat prolonged ALKi sensitivity, whereas untreated HAP cells eventually started re-growing (**Fig. 5E**). A485 produced similar effects (**Fig. S6H**), consistent with prior reports that both HDAC and HAT inhibitors can restrict expression of core regulatory transcription factor genes^50^. These findings indicate that pharmacologic interference with chromatin regulators can suppress the ability of tumor cells to adapt to pharmacological stressors.

Together, these experiments show that phenotypic plasticity is reflected in chromatin accessibility, can be increased or constrained genetically, and can be pharmacologically restrained. By showing that resistance evolves not solely from pre-existing, intrinsically resistant clones but through progressive selection of rare, highly plastic cells, our findings redefine how therapeutic pressure sculpts tumor evolution and point to plasticity itself as a central, targetable driver of resistance.

## DISCUSSION

Across cancers, acquired resistance to targeted therapy has been linked to EMT- and stemness-associated programs, increased metastatic potential, and tolerance to chemotherapy and radiotherapy^51^. Here we show that these recurring features can be understood through a common principle: targeted therapy selects for subpopulations with higher phenotypic plasticity, and this plasticity, rather than a single resistance pathway, drives delayed, multifactorial escape and metastatic competence. We define plasticity as an expanded capacity for therapy-induced state transitions that enables populations to discover adaptive solutions over time. Importantly, this does not necessarily require that highly plastic cells are intrinsically rare at baseline: resistance would be expected to emerge faster when a larger fraction of cells can enter plastic states, and plasticity itself can be induced or amplified by therapeutic and microenvironmental stress.

Our *in silico* model formalized two alternative evolutionary frameworks and predicted that multi-step resistance should progressively enrich subclones with higher plasticity, including when they are scarce at baseline. A key implication was that fitness differences would be largely invisible early during treatment and instead emerge as a delayed expansion of the plastic compartment with continued exposure. Our experiments recapitulated these population-level dynamics: lineage barcoding and single-cell–derived clones revealed a pretreatment fraction with higher adaptation potential that expanded under sustained pressure. Lineage tracing also suggests additional complexity: enrichment profiles were overlapping but not identical across challenges and replicates, consistent with subpopulations that are more capable of adapting to specific stressors, as well as a component of pre-adaptation present before exposure. Here, we focus on plasticity-mediated adaptability across challenges as a general route by which therapeutic pressure can drive resistance evolution. Short-term responses were similar across lineages, but over longer intervals, the high adaptation-potential fraction recovered, adapted to pharmacologic and microenvironmental stressors, and colonized the lung more efficiently *in vivo*.

This selection-for-plasticity framework refines how EMT and stemness should be interpreted in the context of resistance. Rather than serving as universal proximal mechanisms of escape, EMT- and stemness-associated programs mark, and can facilitate, a broader plasticity state that widens the repertoire of phenotypes accessible under selection. In settings where state transitions occur rapidly, plasticity-driven escape may appear as an immediate resistance mechanism, but our data support a time-dependent process in which adaptability emerges through continued selection and reconfiguration.

Consistent with this view, the high adaptation-potential fraction shows increased transcriptional entropy and preferential chromatin accessibility at developmental and cell-identity regulators, features consistent with a regulatory architecture that can be reconfigured under treatment. Importantly, perturbing these identity programs altered long-term outcomes without changing immediate drug sensitivity: partial reprogramming or EMT induction increased adaptation potential in low-plasticity clones, whereas limiting plasticity-sustaining transcription factors reduced resistant outgrowth over longer intervals. Together, these results support plasticity as a convergent, targetable early step on the path to resistance and show that genetic or pharmacologic interventions that constrain plasticity can shift the timing and likelihood of escape.

Although our experiments emphasize non-genetic adaptation, the same principle can extend to settings where resistance is ultimately driven by genetic alterations. By expanding the phenotypic space on which selection operates, plasticity can facilitate reversible reconfiguration and create opportunities for genetic changes that stabilize fitter states. This perspective reconciles drug-specific molecular heterogeneity with recurrent convergence on shared functional properties. Distinct ALK inhibitors drive divergent transcriptional states, yet resistant populations display cross–ALK inhibitor tolerance together with EMT/stemness features, consistent with selection acting on plasticity in addition to drug-specific pathways.

Clinically, progression on targeted therapy is frequently followed by attenuated benefit from subsequent cytotoxic chemotherapy, consistent with resistant disease acquiring broader stress tolerance rather than escaping through a single drug-specific route^52–54^. Resistance also commonly coincides with metastatic progression, and a prevalent interpretation is that this reflects simple arithmetic. Longer disease duration and larger tumor burden provide increased opportunities for dissemination. However, this framing does not explain why resistant disease so often becomes broadly robust across therapies and environments. Our results instead suggest a plausible mechanistic connection: selection for phenotypic plasticity under sustained therapeutic pressure can promote cross-stressor tolerance and increase the likelihood of successful growth in new tissue contexts.

Local microenvironmental constraints are likely to modulate both the emergence and the consequences of this plasticity. Hypoxia, acidosis, nutrient limitation, and paracrine signaling through pathways such as TGF-β and NF-κB can remodel chromatin and redirect transcriptional programs, thereby modulating plasticity and the probability of adaptation^55–57^. Consistent with niche effects, we observe heterogeneity between sites: enrichment patterns differ between primary and metastatic settings, and resistant/high-plasticity cells show superior lung colonization relative to naïve counterparts.

This work focuses on ALK-rearranged NSCLC in long-term culture and xenograft models. Future studies should test how broadly selection for plasticity generalizes across oncogenic drivers, tissues, and immune-competent settings, and should further define quantitative, predictive measures of plasticity in patient samples.

Our findings have translational implications. If treatment selects for plasticity, then intervention timing and combination strategies that constrain state switching may prolong response. In our models, interventions that stabilize cell identity or restrict reprogramming prolonged sensitivity to targeted inhibitors, motivating treatment schedules that intervene during the period when cells begin to reprogram under therapy, before overt resistance emerges. Composite biomarkers of plasticity, including entropy-based metrics, EMT/stemness gene sets, motif-level chromatin signatures, and functional stress-adaptation assays, could stratify risk and guide trials designed to restrain adaptive reprogramming and capture delayed adaptation as an endpoint^58,59^. More broadly, viewing plasticity as a rate-limiting resource provides a framework for integrating non-genetic tolerance with subsequent genetic stabilization of resistant states.

In sum, our data support a model in which therapeutic pressure progressively enriches subpopulations with higher phenotypic plasticity, and this plasticity enables time-dependent adaptation, multifactorial resistance, and metastatic competence. EMT- and stemness-associated programs often accompany, and can potentiate, this capacity but are not, in most contexts, sufficient on their own to confer durable escape. Recognizing plasticity as the trait under selection provides a coherent account of delayed resistance, recurrent transcriptional convergence, and site-specific behavior, and motivates evolution-informed strategies that stabilize identity, modulate paracrine drivers, and constrain adaptive trajectories.

## METHODS

### Cell culture

NSCLC cell line H3122 was obtained from the Lung Cancer Center of Excellence Cell Line depository at Moffitt Cancer Center. H3122 cells were grown in RPMI-1640 containing L-Glutamine (Gibco), supplemented with 10% fetal bovine serum (FBS), 1% penicillin-streptomycin, and 10μg/ml human recombinant insulin (Gibco). Cells were incubated at 37℃ with 5% CO_2_ atmosphere with constant humidity. Cells were passaged as needed using 0.25% trypsin (Gibco). Passage number was kept below 15 passages to maintain consistency in phenotypes.

### Derivation of erLor and erAlec cells

As detailed in Vander Velde (2020)^11^, H3122 cells were subjected to continuous high ALKi concentrations of 2.0uM alectinib or 2.0uM lorlatinib (Fisher Scientific).

### Single-cell Clones derivation

H3122 cells were detached using accutase (Gibco), passed through cell strainers and subjected to sterile flow sorting (BD FACSAria II) to obtain single-cell droplets into six 96-well plates. After careful microscopic evaluation at 1, 3, and 5 days post sorting, we scored 24 wells that had growth from a single cell. Cells were further expanded to proceed with experiments.

### Classification of single-cell derived clones into high, intermediate, or low adaptation potential

We assessed 21 single-cell derived clones’ ability to evolve resistance to alectinib by long-time drug exposure at 500nM (up to 3 months). Among the 21 subclones, 2 showed high adaptation potential, 8 showed intermediate, 11 showed no adaptability scored by their confluency when we stopped the experiment. In order to confirm this preliminary observation and further assess their evolvability of resistance to other TKI’s, duplicate cultures of 10,000 cells/well from each clone seeded into 24-well plates. After 24h, media was replaced with media containing 0.1% DMSO or one ALKi (0.1μM ceritinib (Fisher Scientific), 0.5 μM crizotinib (FisherScientific), 0.5 μM lorlatinib, 0.5 μM alectinib, or 0.1 μM brigatinib (AstaTech). After 40 days of culture, plates were subjected to crystal violet staining and their growth measured by ImageJ by scoring the colony area of wells covered by cells.

### Functional assays

Cells were collected, counted, and plated such that the same number of cells was added to each well of the experimental plate. 24 hours after seeding, media containing the treatment, stressor, or vehicle was added and subsequently replaced every 3-4 days.

Acidosis media was made by adding HCl (6N, VWR) to RPMI-1640 until the pH reached 5.5. Then the acidic media was filtered (0.2um) and the rest of the media components were added. For pseudo-hypoxia media, cobalt chloride hexahydrate (CoCl_2_) was dissolved in PBS then added to RPMI complete media so the final concentration is 200uM. For starvation media, when stressing the evolved resistant lines, the complete media was diluted 1:10 in PBS; when stressing the single-cell derived clones, the complete media was diluted 1:5 in PBS. Cell viability was assessed using crystal violet staining followed by ImageJ quantification or elution, or through Cell Titer Glo assay following manufacturer instructions. Short term assays lasted 3-7 days, long term assays lasted 10-30 days.

### Crystal violet staining

At endpoint, the plate was placed on ice and washed with ice cold PBS (Gibco). Then the cells were fixed with 100% methanol (Fisher Chemical) for 10 minutes on ice. The methanol was aspirated off and the plate was placed at room temperature before 0.5% crystal violet solution in 25% methanol was added for 10 minutes. The crystal violet solution was removed and the plate was submerged in distilled water multiple times to remove excess stain. The plate was placed upside down to dry at room temperature. Once dry, the crystal violet stained plates were imaged on the Amsheran Imager 600 using the colorimetric trans-illumination setting.

### Acetic Acid Elution

After imaging on the Amsherham Imager 600, 10% acetic acid (Fisher Chemical BP2401-500, diluted in double-distilled water) was added to each well, 50ul for a 96-well plate, or 200ul for a 48-well plate. The plate was placed on a shaker for 15 minutes to agitate and elute the crystal violet staining. Then, the acetic acid and crystal violet mixture was diluted with distilled water in a 1:4 ratio. In a new 96-well white-walled clear-bottom plate, 100ul of the solution in each well of the original plate was transferred to a new well in the new plate. The BioTek Synergy Microplate reader read the plate at 590nm. All values were normalized to mean absorbance in DMSO.

### ImageJ quantification

To analyze the image from the Amsherham Imager 600 using ImageJ, the oval selection tool was used to select a single well. The image was then converted to 16-bit. Next, the "Analyze" menu was used to select "Measure" to measure the entire area of the selected well. Then, the "Image" menu was used to select "Adjust" followed by "Threshold." The red mask is used to select empty spaces before selecting “Analyze" and "Measure" again to measure the red area. Subtract the red area from the entire area which was measured first to get the desired measurement.

### Cell Titer Glo Assay

Cell viability was assessed using CellTiterGlo reagent (Promega) following the manufacturers protocol and detected using the luminescence setting on a GloMax luminometer (Promega).

### Cell imaging

Brightfield images were taken on a Zeiss AXIO Observer Z1 microscope to examine cell morphology. For imaging of ex vivo resistant and parental cells, fluorescent channels of 610nm (mCherry) and 509nm (GFP) were used to differentiate between cell lines.

### CloneTracer

H3122 cells were transduced at 30% efficiency (as defined by fluorescence-activated cell sorting analysis of dsRed expression) with CloneTracer neutral DNA barcode library, kindly provided by Frank Stegmeier (Addgene #67267). Barcode-containing cells were selected with puromycin and expanded for 7 days. Barcoded cells were collected and split into conditions. Triplicate cultures of 100,000 cells each were plated into 6-well plates in the presence of 0.1% DMSO, 0.5 μM crizotinib, 0.5 μM alectinib, and 0.5 μM lorlatinib, 0.1 μM ceritinib, 0.1 μM brigatinib, 0.1 μM trametinib, starvation, or acidosis conditions. One aliquot of 100,000 cells was frozen to serve as an initial time point. Cells were collected following 4-8 weeks of cell culture based on their growth in different conditions. Genomic DNA was extracted using proteinase K digest, followed by phenol–chloroform purification. Barcodes were amplified, sequenced, and analyzed following protocols described in (36) and an updated procedure provided on the Addgene website https://www.addgene.org/pooled-library/clontracer. PCR amplification was done with the protocol provided by Stegmier lab, 30 cycles using second-generation ClonTracer primers with the following sequence: 5′-CAA GCA GAA GAC GGC ATA CGA GAT-Variable Sequence-GTG ACT GGA GTT CAG ACG TGT GCT CTT CCG ATC TCT AGC ACT AGC ATA GAG TGC GTA GCT-3′.

### CloneSweeper

H3122 cells were transduced at 5% efficiency, as defined by fluorescence-activated cell sorting analysis of GFP expression, with the neutral DNA barcode library, kindly provided by Shaffer Lab (37). Cells containing barcodes were cultured for 12 days and then divided into treatment groups of 200,000 cells per 10 cm plate. Treatments included DMSO (0.05%), lorlatinib (0.5 µM), alectinib (0.5 µM), or medium adjusted to pH 5.5. An additional aliquot of 200,000 cells was frozen to serve as an initial timepoint. Four plates were prepared for each treatment condition. One plate was harvested and frozen two weeks after treatment initiation to look at intermediate resistance abundance, while the remaining plates were collected once confluent, approximately four weeks for the pH 5.5 condition and five weeks for the lorlatinib and alectinib treatments. Barcode sequences were amplified, sequenced, and analyzed according to the genomic DNA barcode sequencing protocol described in (37).

### Barcode enrichment analysis

For each library, distinct barcode sequences and their corresponding read counts were summarized and merged into a unified count matrix in R (v4.4.0). To account for differences in sequencing depth, barcode counts were normalized to the total number of reads per sample, yielding relative barcode abundances. The resulting normalized and raw count matrices were used for enrichment and comparative evaluations. Barcodes with a frequency above 0.001 in at least 2 samples were kept for further analysis.

### Lentiviral overexpression of Yamanaka factors and Slug

Lentiviral vector for inducible overexpression of *Oct4, Klf4, Myc and Sox2* (OSKM; FUW-tetO-hOKMS) was a gift from Tarjei Mikkelsen (Addgene #51543). Lentiviral vector for inducible expression of *Slug* (Addgene #82162) was Gateway-cloned under a doxycycline induced promoter on a pInducer20 backbone (Addgene #44012). JetPrime reagents and protocol (Polyplus) were followed for transfection into HEK-293T packaging cells. Virus-containing supernatant was collected 24-48h after transfection and filtered through 0.45μm filters (Corning). H3122 low adaptation-potential derivatives were infected with viral lysate containing polybrene (Santa Cruz Biotechnology) before selection in the appropriate antibiotic. To induce expression of either Yamanaka factors or Slug, doxycycline was added to the cell medium for 30 days prior to experiment initiation.

### shRNA knockdown for SOX2

SOX2 inducible lentiviral small hairpin RNA (shRNA) (Tet-pLKO-puro-SOX2) was a gift from Charles Rudin (Addgene #47540). JetPrime reagents and protocol (Polyplus) were followed for transfection into HEK-293T packaging cells. Virus-containing supernatant was collected 24-48h after transfection and filtered through 0.45μm filters (Corning). H3122 high adaptation-potential derivates were infected with viral lysate containing polybrene (Santa Cruz Biotechnology) before selection in the appropriate antibiotic. To induce *SOX2* knockdown, doxycycline was added to the cell medium for 7 days prior to experiment initiation.

### Treatment with Epigenetic Inhibitors

HAP cells were pre-treated with either vehicle (DMSO) or an epigenetic inhibitor A485 (4μM, MedChem Express) or Panobinostat (2 μM, Fisher) for 7 days prior to exposure to ALKi. One arm was co-treated with the epigenetic inhibitor and ALKi to make a total of 3 groups: 1) HAP pretreated with DMSO, then treated with ALKi, ii) HAP pre-treated with epigenetic inhibitor, then treated with ALKi, iii) HAP pre-treated with DMSO, then co-treated with ALKi and epigenetic inhibitor.

### PCR

RNA extraction was performed with RNAeasy (Qiagen) according to the manufacturer’s instructions. Quantitative reverse transcriptase real-time PCR was performed using SensiFAST SYBR Hi-ROX One-Step Kit (Meridian Biosciences, BIO-73005) on a QuantStudio 7 Pro (ThermoFisher). Primer sequences: *NFE2L2* FW, 5’- GAGAGCCCAGTCTTCATTGC-3’; *NFE2L2* RV, 5’-TTGGCTTCTGGACTTGGAAC -3’; *RPL1S* FW, 5’-GATCCGGAAGCTCATCAAAG - 3’, *RPL1S* RV, 5’-GGCTGTACCCTTCCGCTTAC -3’.

### Live Cell Imaging

Cells were plated and allowed to attach for 24 hrs before treatment with either vehicle (DMSO, 0.05%), lorlatinib (0.5μM), or alectinib (0.5μM) along with an epigenetic inhibitor if indicated. Culture medium and drugs were changed every 3 days. Sartorius Incucyte 2020 was used to take images of each well at least every 3 days, and to analyze the images.

#### *In vivo* metastasis subcutaneous

erALKi expressing mCherry and Parental cells expressing GFP or mCherry a 1:1 mixture of erLor:Parental, erAlec:Parental, or Parental (GFP): Parental (mCherry) in RPMI media. Type 3 matrigel was added to each sample in equal volume to the media. The mice were anesthetized with 4% isoflurane for induction, and anesthesia maintenance with 2% isoflurane. 100ul of the cells with matrigel were injected per flank, with 1 million cells per flank. After 4 weeks, both the subcutaneous tumor site and lungs were harvested for flow cytometry analysis. Xenograft studies were performed in accordance with the guidelines of the Institutional Animal Care and All the xenograft studies were performed per the approved procedures of IACUC protocol IS00013344 of the H. Lee Moffitt Cancer Center.

Animals were maintained under AAALAC-accredited specific pathogen-free housing vivarium and veterinary supervision following standard guidelines for temperature and humidity, with a 12-hour light/12-hour dark cycle.

#### *In vivo* metastasis tail vein

erALKi expressing mCherry and Parental cells expressing nuclear GFP or mCherry were prepared. Once counted, a 1:1 mixture of erLor:Parental, erAlec:Parental, or Parental (GFP): Parental (mCherry) in RPMI media. 100ul of the cell-containing media with 100,000 cells per mouse were injected via tail vein. After 4 weeks, the lungs were harvested for flow cytometry analysis.

#### *In vivo* subcutaneous tumors

Aliquots of LAP and HAP cells were prepared with equal volume RPMI media and type 3 matrigel. 375,000 cells were injected per flank. Tumors were allowed to grow until palpable, and alectinib 25mg/kg was delivered daily via gavage. Tumor measurements were taken weekly using electronic calipers. Detection threshold is based on inadequate caliper purchase around lesion. After 19 weeks on alectinib, all three mice bearing HAP-tumors, and one mouse bearing LAP tumors were euthanized and tumors were collected. The remaining 3 mice bearing LAP-tumors were placed on a treatment holiday for 10 weeks until tumors enlarged and were then restarted on alectinib 25mg/kg.

### Tumor digestion and homogenization

Tumors or lungs were digested in RPMI media supplemented with 1 mg/ml collagenase IV (Worthington),1 mg/ml hyaluronidase (Sigma), and 2 mg/ml bovine serum albumin (Fisher Scientific) at 37 °C. After neutralizing with PBS, the tumor solution was filtered with a cell strainer and centrifuged at 1000rpm for 5 minutes. Supernatant was discarded, and cells were either immediately plated for *in vitro* culture, or resuspended in freezing media (40%RPMI complete, 50%FBS, 10%DMSO) and placed in cryovials at - 80 °C.

### Flow cytometry analysis

Tumors or lungs containing GFP/mCherry mixes were digested in RPMI supplemented with 1 mg/ml collagenase IV (Worthington),1 mg/ml hyaluronidase (Sigma), and 2 mg/ml bovine serum albumin (Fisher Scientific) at 37 °C. Pellets were suspended in PBS and 0.1 μg/ml DAPI (Sigma). Percent of population that is mCherry+ and GFP+ was determined using a MACSQuant Flow Cytometer.

### Plasma luciferase

Gateway cloning (Invitrogen) was used to generate plasmids with pLenti 6.3 backbone (Invitrogen) and C-luciferase (ThermoFisher, cat:16150). JetPrime reagents and protocol (Polyplus) were followed for transfection into HEK-293T packaging cells. Virus-containing supernatant was collected 24-48h after transfection and filtered through 0.45μm filters (Corning). H3122 cells were infected with viral lysate containing polybrene (Santa Cruz Biotechnology) before selection in the appropriate antibiotic. After selection, luciferase expression was checked by collecting 200ul of the conditioned media after 6 hours. The C-luciferase assay was then conducted.

#### *In vivo* luciferase monitoring

LAP, HAP, or Parental cells secreting C-luciferase were prepared for injection such that 100,000 cells were injected via tail vein per mouse. Every two weeks, blood from the tail vein of each mouse was collected into ethylenediaminetetraacetic acid (EDTA) -coated tubes (ThermoScientific) and centrifuged at 2000g for 10 minutes. The plasma was collected and diluted at 1:100 with PBS before the C-luciferase assay was conducted.

### C-Luciferase Assay

Vargulin, cyprindina luciferase (NanoLight Technologies), was prepared in 0.5mg/ml stock in acidified methanol (5ul of 6N HCL per 1ml methanol). A 1:1000 dilution of stock luciferase was prepared in PBS to make working concentrations. In triplicate, 20ul of the conditioned media or diluted plasma sample was plated in a white-walled 96-well plate. The assay was run with 100ul of luciferase added per well with an integration time of 2 seconds.

### Bulk RNA-seq

Duplicate cultures of H3122 parental cells and single-cell derived clones were cultured until 60-70% confluent, then lysed directly by adding RNA lysis buffer. RNA was isolated for parental cells and single-cell clones using an RNAEasy Minikit (Qiagen). Reads were generated using a Novaseq illumina platform at 30,000,000 reads per sample. Alignment was achieved using HiSat2 and the human hg19 reference genome. Normalized reads were obtained from DeSeq2. Additional unsupervised clustering and PCA analysis was performed using R.

### Gene set enrichment analysis

GSEA was used to determine the gene sets enriched in high, intermediate and low evolvable clones versus parental cell line RNA-seq profiles. We used the MSigDB Hallmark [PMID: 26771021] as the predefined gene sets and performed 10,000 permutations by gene set to determine the p-values. Gene sets with false discovery rate q-value ≤0.25 were considered as significantly enriched.

### Single cell transcriptome analysis

Data was originally reported in Vander Velde (2020)^11^, briefly, cells were exposed to either no treatment (naive), or to 112 days of lorlatinib or alectinib. Approximately 1,000 cells were collected per sample and processed using the 10X Genomics pipeline on the Illumina NextSeq 500 instrument. In this article, the data has been further analyzed by Gini Score.

### Gini Analysis

Single cell transcriptomic data was analyzed using the “DescTools” and “Seurat” packages on R.

### ATAC-seq

Duplicate cultures of H3122 parental cells and cell-line derivatives (two clones from each category such as high, intermediate and low adaptability potential clones, and two independently derived erLor and erAlec replicates) were cultured in complete RPMI until 60-70% confluent. Cells were collected and frozen. ATAC-sequencing was performed by ActiveMotif for at least 50,000 reads. Usable reads were aligned to the hg38 reference genome. Bioinformatic analysis was done in collaboration with Berkley Gryder using a previously established pipeline which can be found at https://github.com/GryderLab/gryderlab_pipeline. Briefly, ATAC-seq reads were matched with bulk RNA-seq data from the same clone. Motif enrichment analysis was performed using HOMER and biological processes were identified using ToppGene. Biological processes with p-value less than 1×10^-6^ were considered significant.

### Computational Model

The agent-based model (ABM) includes three cell types—sensitive, baseline-plasticity persisters, and high-plasticity persisters—each capable of proliferation and death. Division and death rates depend on treatment status and local environmental conditions (Supplementary Methods). Mutations arise stochastically at division and are assumed beneficial, providing incremental fitness gains toward resistance. Mutations are mechanism-agnostic (genetic or epigenetic) and occur only upon division. Treatment is modeled as a binary, domain-wide perturbation: sensitive cells are fully inhibited and rapidly decline, while persister populations maintain net-zero growth until resistance mutations appear.

The ABM is implemented on a 10,000-site two-dimensional grid. Agents interact with their local environment through a Moore neighborhood of eight adjacent sites. Initial populations are randomly seeded according to experimentally or theoretically motivated proportions. At each discrete time step, agents update asynchronously following probabilistic rules for division, death, and mutation. Fifty independent stochastic realizations were run to capture variability.

To complement the ABM and isolate the effects of stochasticity and spatial structure, a deterministic ODE model was developed under identical biological assumptions and rate parameters, modeling population-level dynamics with a carrying capacity of 1,000 cells. Python was used for parameter estimation, data processing, visualization, and to implement both the ABM and ODE models. The ODE simulations, being deterministic, were run once per parameter set. All simulation scripts, parameter files, and analysis notebooks are publicly available at https://github.com/mfroid/plasticityModels/. All ABM data is available upon request from the authors. See **Supplemental Methods** for additional information.

### Statistical Analyses

Statistical analysis was performed with Prism 10 (GraphPad) or R statistical software. Details of statistical tests are provided in Supplementary Table 1. \**p*<0.05, \*\**p*<0.01, \*\*\**p*<0.001, and \*\*\*\**p*<0.0001.

## Supporting information

Supplementary Methods_Math Model

Table of Statistics

## DATA AND CODE AVAILABILITY

- All simulation scripts, parameter files, and analysis notebooks are publicly available at https://github.com/mfroid/plasticityModels/. All ABM data is available upon request from the authors.
- ATAC-seq data is publicly available at https://www.ncbi.nlm.nih.gov/sra/PRJNA1400308. The previously established analysis pipeline can be found at https://github.com/GryderLab/gryderlab_pipeline.
- CloneTracer and CloneSweeper datasets can be found at https://www.ncbi.nlm.nih.gov/sra/PRJNA1400309.
- Single Cell RNA sequencing data was previously reported in (6) and can be found at https://www.ncbi.nlm.nih.gov/geo/query/acc.cgi?acc=GSE144282.
- Bulk RNA sequencing data can be found at https://www.ncbi.nlm.nih.gov/sra/PRJNA1400309.
- All data reported in this paper are available from the corresponding author upon request.

## AUTHOR CONTRIBUTIONS

AM conceived of the project. AB and XA designed and performed wet-lab experiments and data analysis. MF, RBC, and JM, designed and performed *in silico* experiments under the supervision of AM, JC, and DB. JR performed growth rate assays under the supervision of AB and AM. XA generated the bulk RNA-seq samples and performed the analysis. XA performed the Clone Tracer experiment and analyzed it with VT. RV contributed to making the CloneSweeper technology and liaised our use of it. AB performed the CloneSweeper experiment. AB generated the ATAC-seq samples and performed the analysis under the supervision of BG. AB, VT, and AM designed the schematics. AB, VT, and AM wrote the manuscript. JC, DB, AA, and VT made substantial input to the direction, design, and execution of the project. All authors reviewed the manuscript.

## ACKNOWLEDGEMENTS

We thank M. Henry for their contribution to the animal work. This work has been supported in part by the Flow Cytometry Core and the Molecular Genomics Core at the H. Lee Moffitt Cancer Center and Research Institute.

## REFERENCES

1. Strebhardt, K. & Ullrich, A. Paul Ehrlich’s magic bullet concept: 100 years of progress. Nat. Rev. Cancer 8, 473–480 (2008).

2. Reck, M. & Rabe, K. F. Precision Diagnosis and Treatment for Advanced Non–Small-Cell Lung Cancer. N. Engl. J. Med. 377, 849–861 (2017).

3. Huang, L., Jiang, S. & Shi, Y. Tyrosine kinase inhibitors for solid tumors in the past 20 years (2001–2020). J. Hematol. Oncol.J Hematol Oncol 13, 143 (2020).

4. Lung Cancer Survival Rates | 5-Year Survival Rates for Lung Cancer. American Cancer Society https://www.cancer.org/cancer/types/lung-cancer/detection-diagnosis-staging/survival-rates.html.

5. Ramos, P. & Bentires-Alj, M. Mechanism-based cancer therapy: resistance to therapy, therapy for resistance. Oncogene 34, 3617–3626 (2015).

6. Shaffer, S. M. et al. Rare cell variability and drug-induced reprogramming as a mode of cancer drug resistance. Nature 546, 431–435 (2017).

7. Marine, J.-C., Dawson, S.-J. & Dawson, M. A. Non-genetic mechanisms of therapeutic resistance in cancer. Nat. Rev. Cancer 20, 743–756 (2020).

8. Sharma, S. V. et al. A Chromatin-Mediated Reversible Drug-Tolerant State in Cancer Cell Subpopulations. Cell 141, 69–80 (2010).

9. Nguyen, A., Yoshida, M., Goodarzi, H. & Tavazoie, S. F. Highly variable cancer subpopulations that exhibit enhanced transcriptome variability and metastatic fitness. Nat. Commun. 7, 11246 (2016).

10. Pérez-González, A., Bévant, K. & Blanpain, C. Cancer cell plasticity during tumor progression, metastasis and response to therapy. Nat. Cancer 4, 1063–1082 (2023).

11. Vander Velde, R., et al. Resistance to targeted therapies as a multifactorial, gradual adaptation to inhibitor specific selective pressures. Nat. Commun. 11, 2393 (2020).

12. França, G. S. et al. Cellular adaptation to cancer therapy along a resistance continuum. Nature 631, 876–883 (2024).

13. Scarborough, J. A., et al. Identifying States of Collateral Sensitivity during the Evolution of Therapeutic Resistance in Ewing’s Sarcoma. iScience 23, (2020).

14. Fukuda, K. et al. Epithelial-to-Mesenchymal Transition Is a Mechanism of ALK Inhibitor Resistance in Lung Cancer Independent of ALK Mutation Status. Cancer Res. 79, 1658–1670 (2019).

15. Deng, S. et al. Ectopic JAK–STAT activation enables the transition to a stem-like and multilineage state conferring AR-targeted therapy resistance. Nat. Cancer 3, 1071–1087 (2022).

16. Byers, L. A. et al. An Epithelial–Mesenchymal Transition Gene Signature Predicts Resistance to EGFR and PI3K Inhibitors and Identifies Axl as a Therapeutic Target for Overcoming EGFR Inhibitor Resistance. Clin. Cancer Res. 19, 279–290 (2013).

17. Zhang, Z. et al. Activation of the AXL kinase causes resistance to EGFR-targeted therapy in lung cancer. Nat. Genet. 44, 852–860 (2012).

18. Holohan, C., Van Schaeybroeck, S., Longley, D. B. & Johnston, P. G. Cancer drug resistance: an evolving paradigm. Nat. Rev. Cancer 13, 714–726 (2013).

19. Jiang, C. et al. Innate immunity and the NF-κB pathway control prostate stem cell plasticity, reprogramming and tumor initiation. Nat. Cancer 1–22 (2025) doi:10.1038/s43018-025-00994-3.

20. Cammareri, P. et al. Loss of colonic fidelity enables multilineage plasticity and metastasis. Nature 1–10 (2025) doi:10.1038/s41586-025-09125-5.

21. Lee, J. H. et al. TGF-β and RAS jointly unmask primed enhancers to drive metastasis. Cell 187, 6182–6199.e29 (2024).

22. Xu, J., Lamouille, S. & Derynck, R. TGF-β-induced epithelial to mesenchymal transition. Cell Res. 19, 156–172 (2009).

23. Yeh, H.-W. et al. PSPC1 mediates TGF-β1 autocrine signalling and Smad2/3 target switching to promote EMT, stemness and metastasis. Nat. Cell Biol. 20, 479–491 (2018).

24. Kim, B. N. et al. TGF-β induced EMT and stemness characteristics are associated with epigenetic regulation in lung cancer. Sci. Rep. 10, 10597 (2020).

25. Su, J. et al. TGF-β orchestrates fibrogenic and developmental EMTs via the RAS effector RREB1. Nature 577, 566–571 (2020).

26. O’Hagan, S., Wright Muelas, M., Day, P. J., Lundberg, E. & Kell, D. B. GeneGini: Assessment via the Gini Coefficient of Reference “Housekeeping” Genes and Diverse Human Transporter Expression Profiles. Cell Syst. 6, 230–244.e1 (2018).

27. Torre, E. et al. Rare Cell Detection by Single-Cell RNA Sequencing as Guided by Single-Molecule RNA FISH. Cell Syst. 6, 171–179.e5 (2018).

28. Estrella, V. et al. Acidity Generated by the Tumor Microenvironment Drives Local Invasion. Cancer Res. 73, 1524–1535 (2013).

29. Pavlova, N. N., Zhu, J. & Thompson, C. B. The hallmarks of cancer metabolism: Still emerging. Cell Metab. 34, 355–377 (2022).

30. Gilmartin, A. G., et al. GSK1120212 (JTP-74057) Is an Inhibitor of MEK Activity and Activation with Favorable Pharmacokinetic Properties for Sustained In Vivo Pathway Inhibition. Clin. Cancer Res. 17, 989–1000 (2011).

31. Blumenschein, G. R. et al. A randomized phase II study of the MEK1/MEK2 inhibitor trametinib (GSK1120212) compared with docetaxel in KRAS-mutant advanced non-small-cell lung cancer (NSCLC). Ann. Oncol. 26, 894–901 (2015).

32. Verstovsek, S. et al. Safety and Efficacy of INCB018424, a JAK1 and JAK2 Inhibitor, in Myelofibrosis. N. Engl. J. Med. 363, 1117–1127 (2010).

33. Li, J., Kim, S. G. & Blenis, J. Rapamycin: One Drug, Many Effects. Cell Metab. 19, 373– 379 (2014).

34. Lambert, A. W., Zhang, Y. & Weinberg, R. A. Cell-intrinsic and microenvironmental determinants of metastatic colonization. Nat. Cell Biol. 26, 687–697 (2024).

35. Combined Use of Tail Vein Metastasis Assays and Real-Time In Vivo Imaging to Quantify Breast Cancer Metastatic Colonization and Burden in the Lungs. https://app.jove.com/v/60687/combined-use-tail-vein-metastasis-assays-real-time-vivo-imaging-to.

36. Bhang, H. C. et al. Studying clonal dynamics in response to cancer therapy using high-complexity barcoding. Nat. Med. 21, 440–448 (2015).

37. Velde, R. J. V., Ng, R. W. S., Coté, C. & Shaffer, S. M. Multiplexed enrichment and tracking of lineages with CloneSweeper. 2026.01.30.700779 Preprint at 10.64898/2026.01.30.700779 (2026).

38. Charles, J. P. et al. Monitoring the dynamics of clonal tumour evolution in vivo using secreted luciferases. Nat. Commun. 5, 3981 (2014).

39. Yue, D. et al. Hedgehog/Gli promotes epithelial-mesenchymal transition in lung squamous cell carcinomas. J. Exp. Clin. Cancer Res. 33, 34 (2014).

40. Li, D. et al. Chromatin Accessibility Dynamics during iPSC Reprogramming. Cell Stem Cell 21, 819–833.e6 (2017).

41. Buenrostro, J. D., Giresi, P. G., Zaba, L. C., Chang, H. Y. & Greenleaf, W. J. Transposition of native chromatin for fast and sensitive epigenomic profiling of open chromatin, DNA-binding proteins and nucleosome position. Nat. Methods 10, 1213–1218 (2013).

42. Takahashi, K. et al. Induction of Pluripotent Stem Cells from Adult Human Fibroblasts by Defined Factors. Cell 131, 861–872 (2007).

43. Chang, T.-H. et al. Slug Confers Resistance to the Epidermal Growth Factor Receptor Tyrosine Kinase Inhibitor. Am. J. Respir. Crit. Care Med. 183, 1071–1079 (2011).

44. Guo, W. et al. Slug and Sox9 Cooperatively Determine the Mammary Stem Cell State. Cell 148, 1015–1028 (2012).

45. Sun, Y., Song, G.-D., Sun, N., Chen, J.-Q. & Yang, S.-S. Slug overexpression induces stemness and promotes hepatocellular carcinoma cell invasion and metastasis. Oncol. Lett. 7, 1936–1940 (2014).

46. Moon, J. H., Lee, S. H. & Lim, Y. C. Wnt/β-catenin/Slug pathway contributes to tumor invasion and lymph node metastasis in head and neck squamous cell carcinoma. Clin. Exp. Metastasis 38, 163–174 (2021).

47. Qian, D. Z., et al. Targeting Tumor Angiogenesis with Histone Deacetylase Inhibitors: the Hydroxamic Acid Derivative LBH589. Clin. Cancer Res. 12, 634–642 (2006).

48. Ellis, L. et al. Histone deacetylase inhibitor panobinostat induces clinical responses with associated alterations in gene expression profiles in cutaneous T-cell lymphoma. Clin. Cancer Res. Off. J. Am. Assoc. Cancer Res. 14, 4500–4510 (2008).

49. Shendy, N. A. M. et al. Group 3 medulloblastoma transcriptional networks collapse under domain specific EP300/CBP inhibition. Nat. Commun. 15, 3483 (2024).

50. Gryder, B. E. et al. Histone hyperacetylation disrupts core gene regulatory architecture in rhabdomyosarcoma. Nat. Genet. 51, 1714–1722 (2019).

51. Gupta, P. B., Pastushenko, I., Skibinski, A., Blanpain, C. & Kuperwasser, C. Phenotypic Plasticity: Driver of Cancer Initiation, Progression, and Therapy Resistance. Cell Stem Cell 24, 65–78 (2019).

52. Zeng, Z. et al. Reduced chemotherapy sensitivity in EGFR-mutant lung cancer patient with frontline EGFR tyrosine kinase inhibitor. Lung Cancer 86, 219–224 (2014).

53. Xu, J. et al. Comparison of outcomes of tyrosine kinase inhibitor in first- or second-line therapy for advanced non-small-cell lung cancer patients with sensitive EGFR mutations. Oncotarget 7, 68442–68448 (2016).

54. Loria, R. et al. Cross-Resistance Among Sequential Cancer Therapeutics: An Emerging Issue. Front. Oncol. 12, (2022).

55. Palma, F. R. et al. Histone H3.1 is a chromatin-embedded redox sensor triggered by tumor cells developing adaptive phenotypic plasticity and multidrug resistance. Cell Rep. 43, 113897 (2024).

56. Chakraborty, A. A. et al. Histone demethylase KDM6A directly senses oxygen to control chromatin and cell fate. Science 363, 1217–1222 (2019).

57. Rolver, M. G., et al. Tumor microenvironment acidosis favors pancreatic cancer stem cell properties and in vivo metastasis. iScience 28, (2025).

58. Zhang, J., Cunningham, J. J., Brown, J. S. & Gatenby, R. A. Integrating evolutionary dynamics into treatment of metastatic castrate-resistant prostate cancer. Nat. Commun. 8, 1816 (2017).

59. Smalley, I. et al. Leveraging transcriptional dynamics to improve BRAF inhibitor responses in melanoma. EBioMedicine 48, 178–190 (2019).

