## Supplementary Figures for "Selection for targeted therapy resistance leads to an indirect selection for higher phenotypic plasticity and enhanced evolvability to orthogonal stressors"

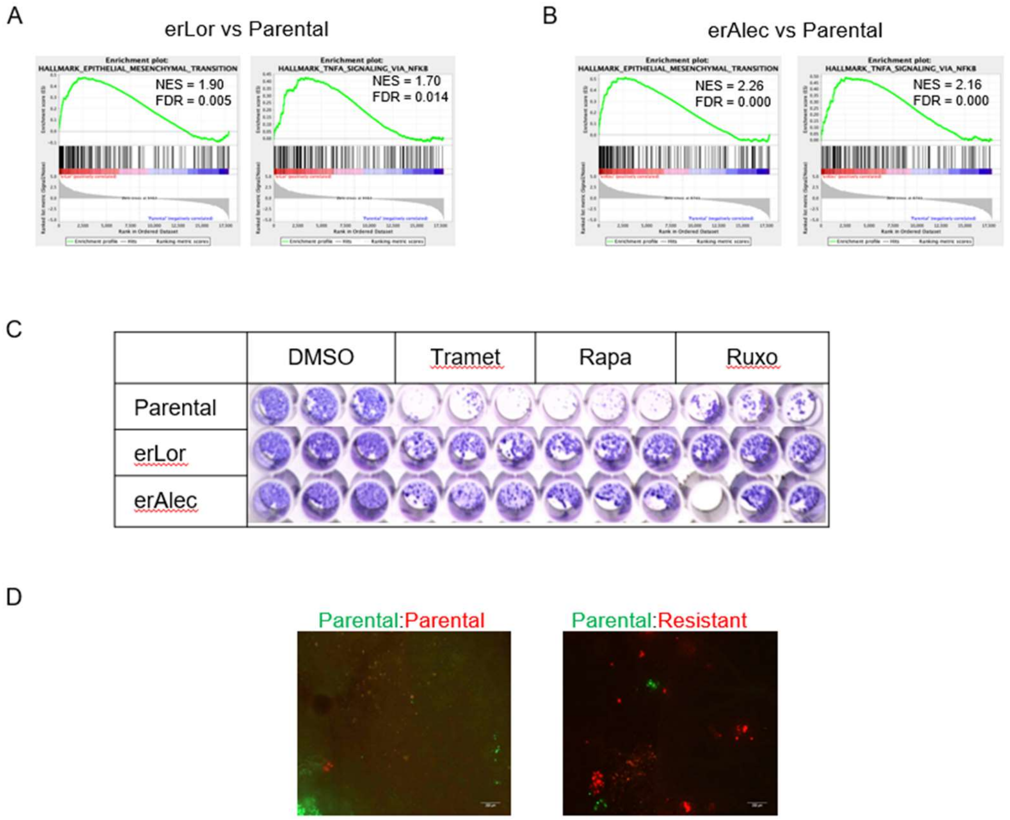


**Supplementary Figure 1. A.** Enrichment plots from GSEA of hallmark gene sets epithelial-to-mesenchymal (EMT) and tumor necrosis factor-alpha (TNFa) via nuclear factor kappa B (NF-κB) for erLor versus Parental cells or (**B**) erAlec vs Parental cells from RNA-sequencing. NES and false discovery rate (FDR) shown. **C.** Crystal violet staining after 15 days of Parental, erLor, or erAlec cells in given pharmacological stress condition relating to Fig. 1H. **D.** Fluorescent images ex vivo from experiment described in Fig. 1J.

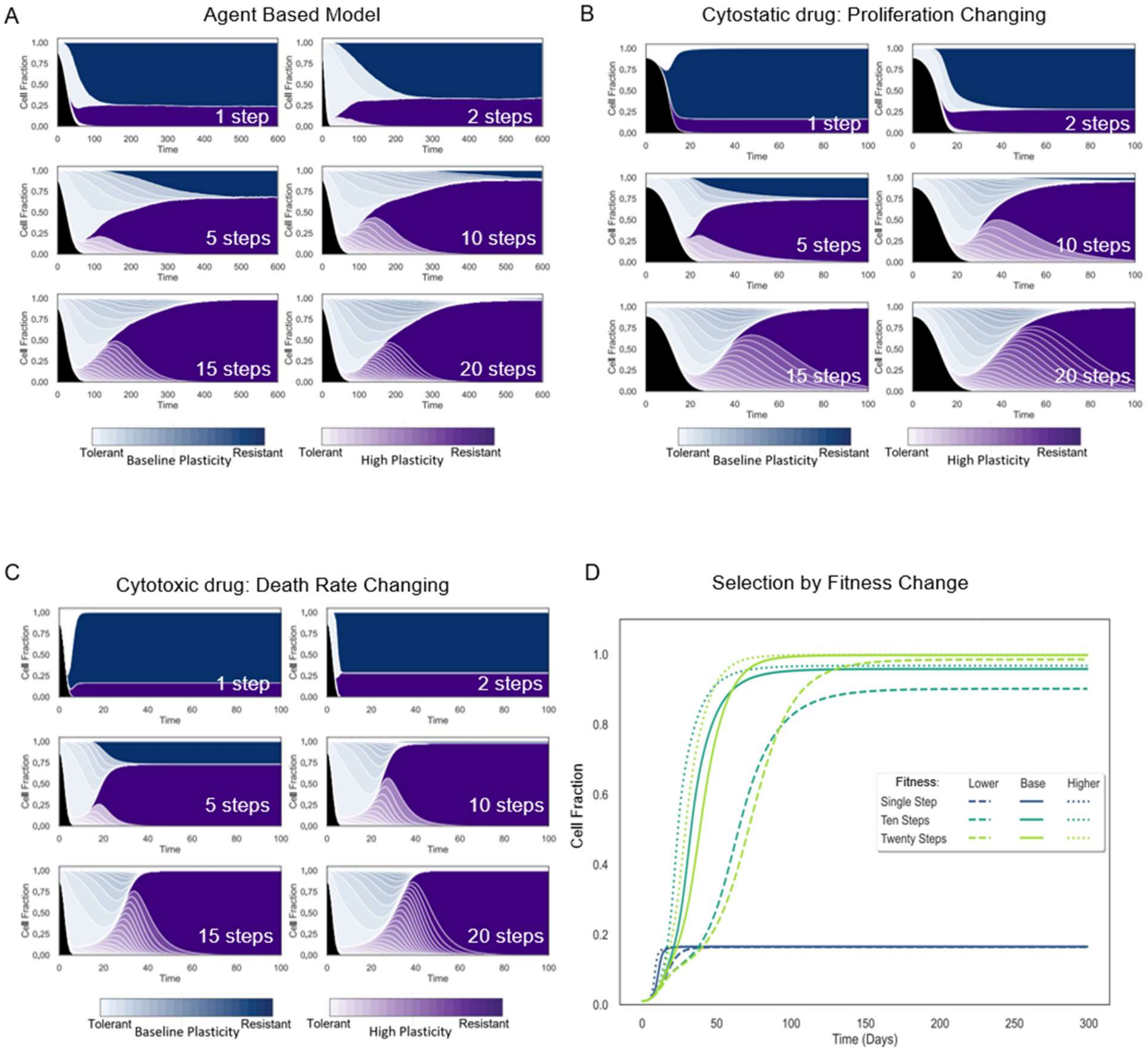


**Supplementary Figure 2. A.** ABM model results. Cell fraction of each population over 600 timesteps based on the number of steps in each simulation shown. **B**. ODE model with proliferation rate ($G_s$) varied to model cytostatic drug effects. **C.** ODE model with death rate ($D$) varied to model cytotoxic drug effects. **D.** Simulated growth curve with differing fitness values in ODE model based on number of steps defined.

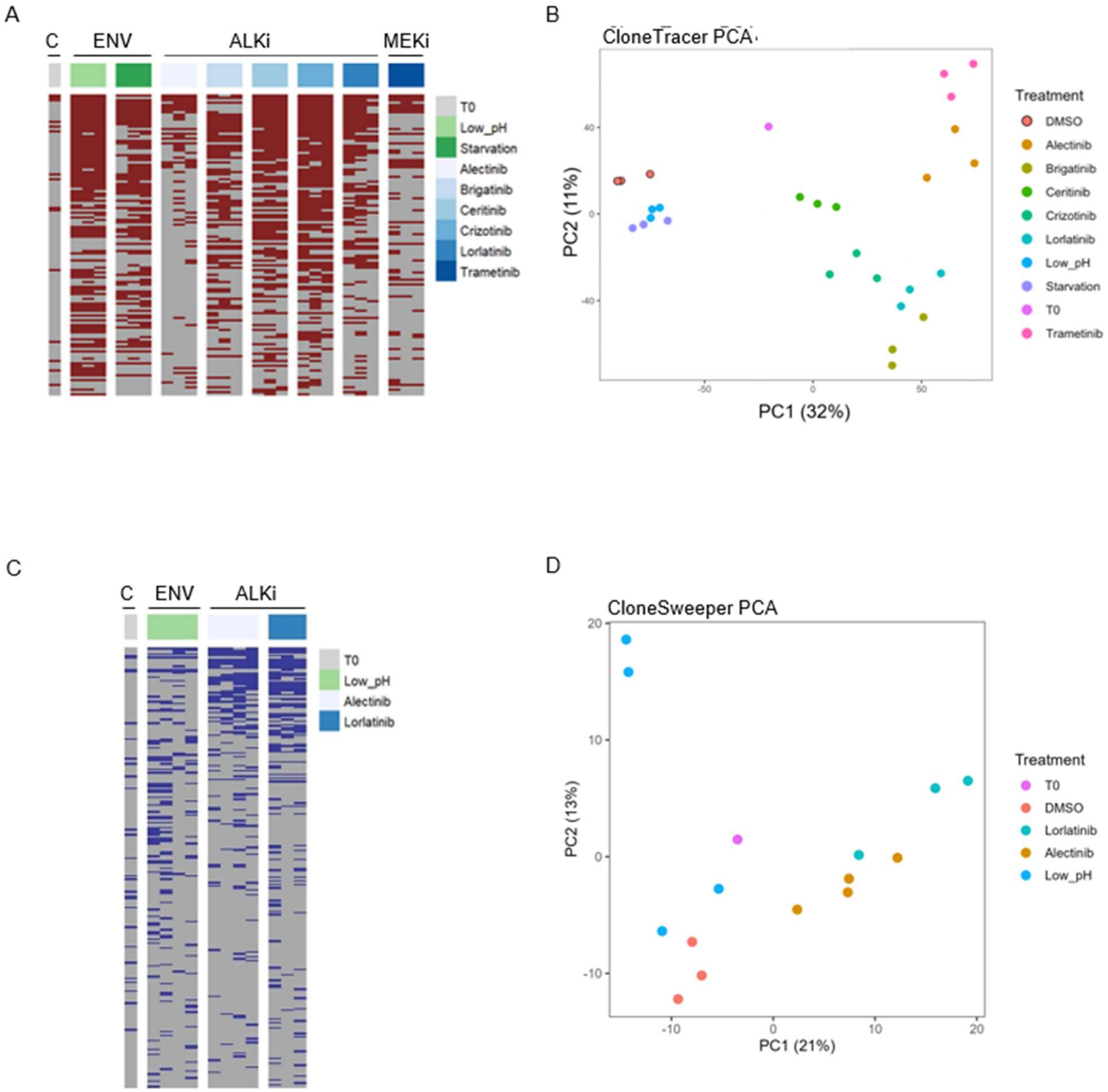


**Supplementary Figure 3. A.** Heatmap of top 20% of barcodes ranked by number of samples in which it is present. Each row represents a given barcode. Each column represents a given replicate of a condition, (C = control, T0= time zero, initial time point; ENV = environmental stressor, ALKi= ALK inhibitor, MEKi= MEK inhibitor). **B.** Principal component analysis (PCA) from barcode analysis of CloneTracer lineage tracing. **C.** CloneSweeper barcodes present in >0.1% of population. Arranged in descending order. **D.** PCA of CloneSweeper data.

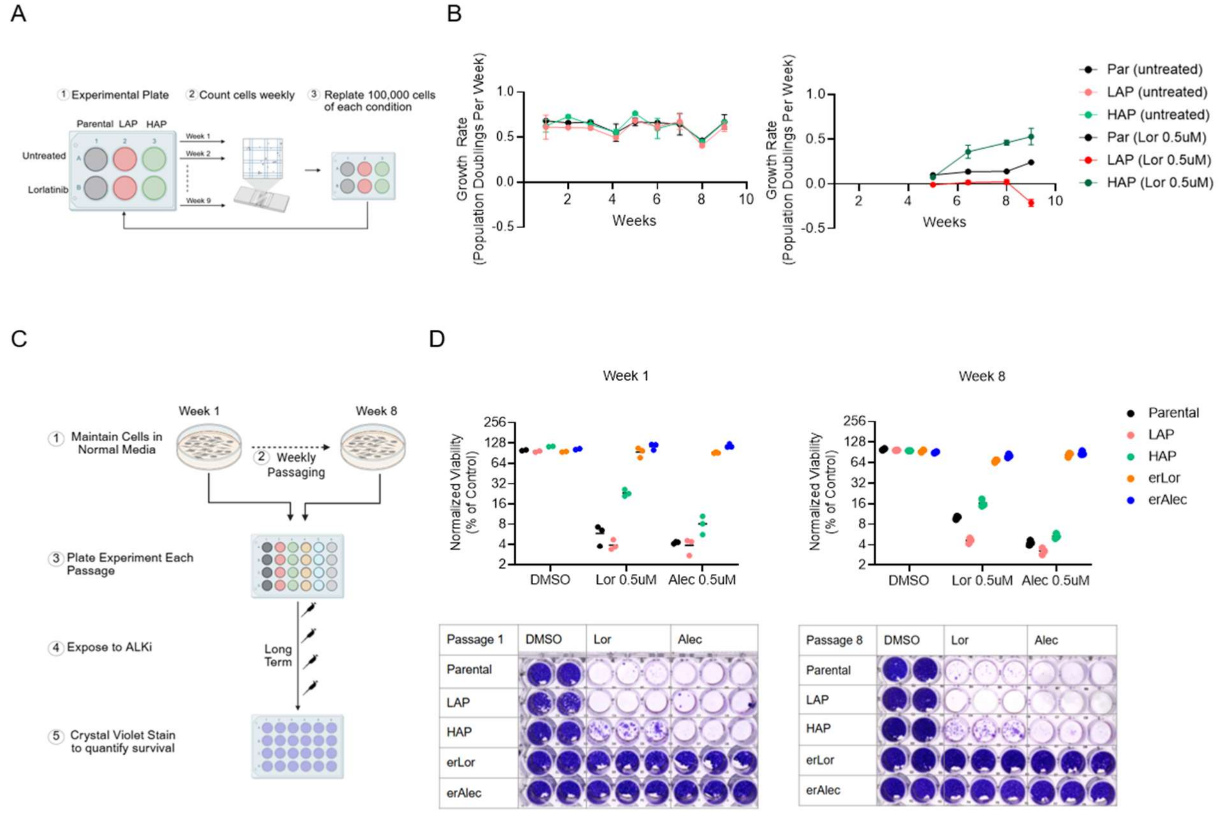


**Supplementary Figure 4. A.** Schematic of growth rate assay. **B.** Growth rate traces from (left) untreated or (right) treated Parental, LAP, and HAP cell lines. Treated conditions had too few cells to warrant collection and counting in weeks 1-5. **C.** Schematic of phenotypic stability assay. **D.** Quantification (top) of crystal violet staining (bottom) after 4 weeks in treatment following one passage (left) or eight passages (right). Two-way ANOVA with Tukey's comparison to Parental cells in Supplementary Table 1.

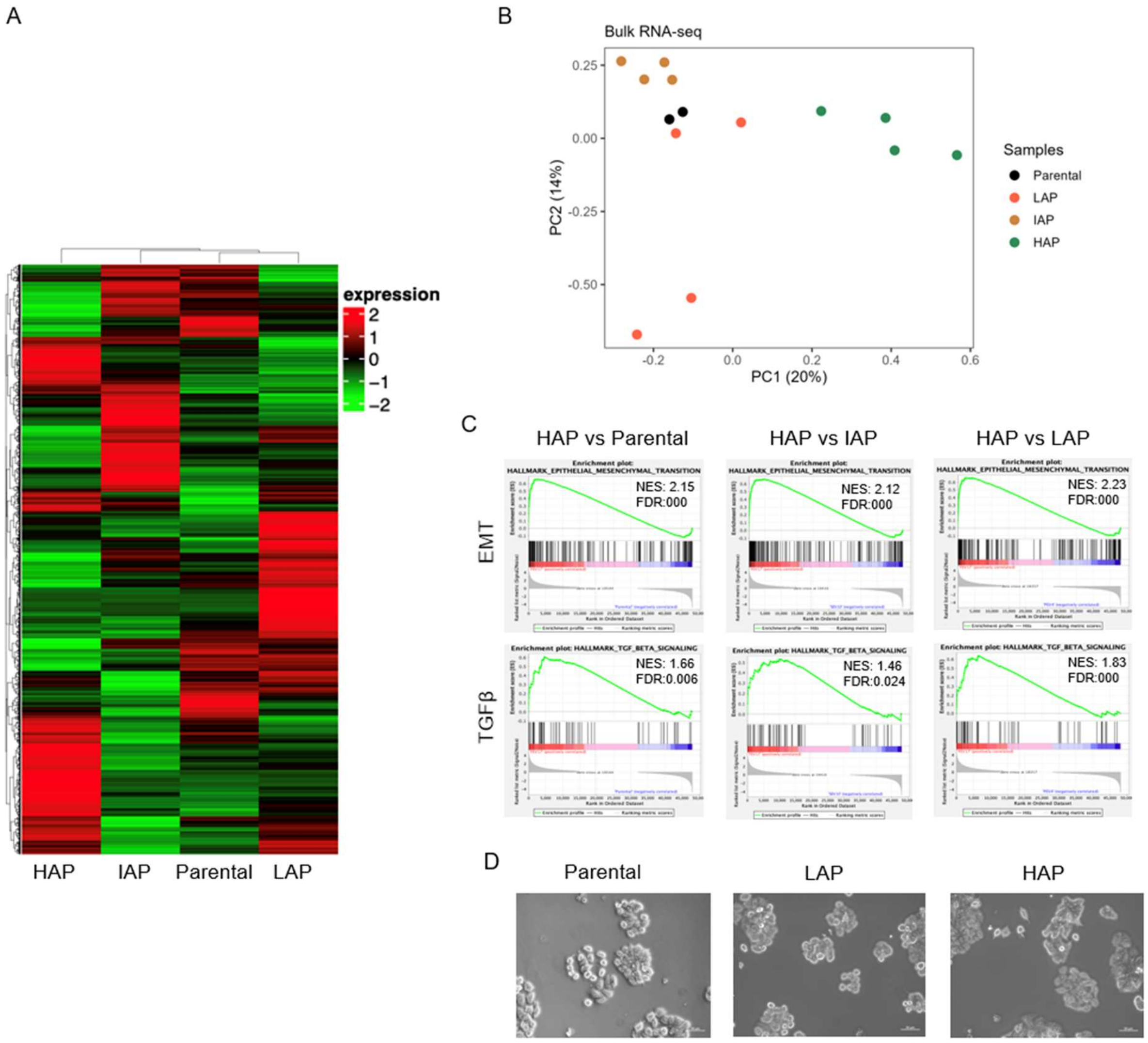


**Supplementary Figure 5. A**. Unsupervised hierarchical clustering of gene expression between Parental, HAP, IAP, or LAP cells. **B**. PCA plot of bulk RNA sequencing. **C.** Enrichment plots from GSEA of hallmark gene sets EMT and transforming growth factor-beta (TGFb) for HAP versus Parental, HAP versus IAP, or HAP versus LAP cells from bulk RNA-sequencing. **D.** Microscopy photos of Parental, LAP and HAP cells at 20x magnification (Scale bar 50μm).

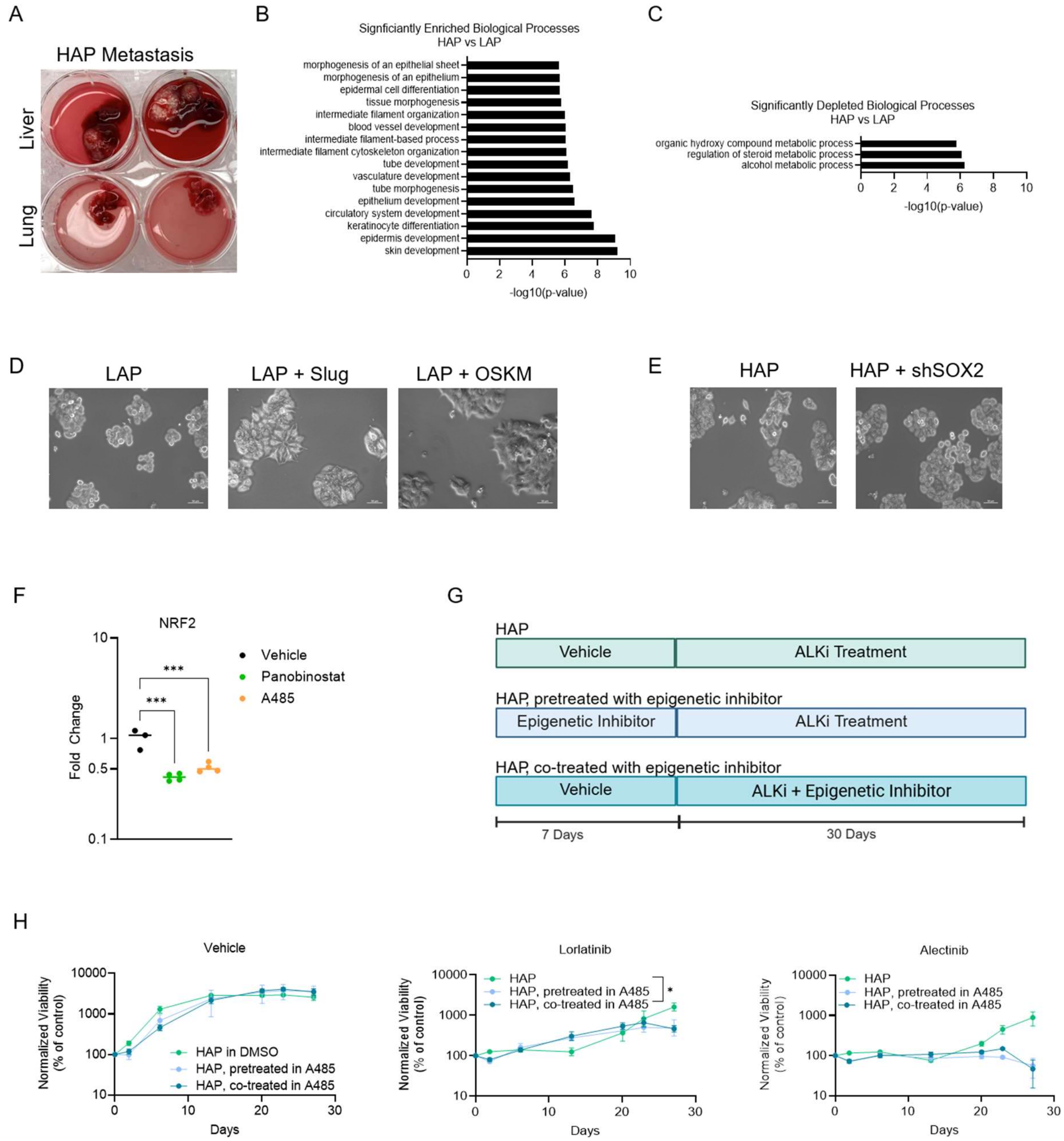


**Supplementary Figure 6. A.** Images of livers with metastasis and lungs from two of the mice injected with HAP cells, at endpoint. **B.** Significantly enriched or (**C**) significantly depleted biological process from bulk ATAC-sequencing analysis of HAP versus LAP cells. –log10(p-value) shown. **D.** Representative bright-field images of LAP, LAP+Slug, and LAP +OSKM cells at 20x magnification (Scale bar 50µm). **E.** Representative bright-field images of HAP and HAP + shSOX2 cells at 20x magnification (Scale bar 50µm). **F.** Quantification of reverse transcriptase real time PCR of NRF2 after 7 days in treatment with epigenetic inhibitor. Ordinary One-way ANOVA with Dunnett's multiple comparison test shown. **G.**

Experimental schematic for treatment with epigenetic and ALK inhibition. **H.** Quantification by live cell imaging of percentage of area covered by cells treated with vehicle (DMSO, 0.025%) (left), lorlatinib 0.5μM (middle), or alectinib 0.5μM (right) with pre-treatment of co-treatment with A485 (4μM). Two-way ANOVA with repeated measurements and Tukey's multiple comparison used; comparison to HAP shown.
