## Supplementary Methods_Math Model for "Selection for targeted therapy resistance leads to an indirect selection for higher phenotypic plasticity and enhanced evolvability to orthogonal stressors"


### Summary

To test the validity of our hypothesis we developed two mathematical models, one is an ordinary differential equation model (ODE) and the other a spatial agent-based model (ABM). The ODE starts with a population of 10000 cells, all of which are allowed to divide, die and mutate according to a system of ordinary differential equations. For the ABM, 10000 initial cells are placed on a 2D lattice and allowed to divide, die, and mutate according to a set of simple rules (see end of document for further details). Whenever a cell probabilistically acquires a new molecular change (the model is agnostic as to whether this is a genetic or epigenetic change) it becomes a discrete step closer to becoming fully resistant. Of the 10000 initial cells in both the ODE and ABM, a small subset of the cells begin the simulation in a "persister state" exhibiting net zero growth under treatment, which represents a cell state that is tolerant of treatment but cannot expand under treatment. Of these persister state cells, some are endowed with 2x higher plasticity and therefore have a 2x higher probability of acquiring new changes. We chose to begin simulations with the ratio of cells with base line plasticity to cells with increased plasticity at a ratio of 1:10. The higher plasticity cells are 2x more evolvable based on the HAP frequency in our experiments and the observation that the higher fitness potential of these cells only becomes available over longer treatment courses.

By keeping the time interval from initial placement of cells to when the development of overt resistance becomes fixed, this model allowed us to explore the population dynamics associated with either single-step or multi-step resistance and ask which of this mechanism better explains experimental observations.

We found that multi-step resistance (5 or more discrete steps) leads to the selection and expansion of clones with higher evolvability, even if these cells represent a minority of the starting population (1:10 and 1:100). However, in the single-step or two-steps resistance scenarios, plastic cells remain rarer throughout the whole observation period, and, as a result of a numbers game, resistance reproducibly (across replicate simulations) emerges from the more frequent cells with a base level plasticity.

Additionally, we found differences in the time scale in which resistance develops between the ABM and the ODE models. This appeared to be due to the competition for space in the ABM, as all parameters and rates were shared between the two models.


### Model overview

The purpose of both the ordinary differential equation (ODE) model and the agent-based model (ABM) is to study the evolution of cancer in response to treatment. More specifically, we are using these models to explore the impact of multiple steps towards resistance in the same time as a single step towards resistance, and if there is increased selection for more plastic cells as the number of steps towards resistance increases. The decision to use two different modeling paradigms was due to the computational demands that an ABM necessitates. The ODE, being continuous and deterministic, requires much less computational power and served as a tool to test hypotheses before implementing the ABM. The decision to use an ABM was to explore how spatial constraints and the competition for space, much like those found in realistic living systems, impacts selection for more plastic cells.

**Model assumptions**

The models includes three initial cell types: sensitive cells, persister cells with baseline plasticity, and persister cells with high plasticity. All cell types are able to divide and die, with the rate of division and death determined by various factors such as treatment and environmental conditions. Sensitive cells are completely sensitive to treatment and die quickly, while persister cells have net-zero growth rates initially. However, persister cells can gain mutations that allow them to evolve towards resistance. Mutations only occur during cell division, and high plasticity cells are able to evolve twice as fast as baseline plasticity cells. For both models, mutations occur stochastically during cell division and are assumed to be beneficial, conferring incremental fitness advantages toward resistance. Baseline and High plasticity cells differ in their rates of evolutionary adaptation, with high plasticity cells acquiring beneficial mutations at twice the rate of baseline cells. Treatment is modeled after tyrosine kinase inhibitors commonly used to treat non-small cell lung carcinoma. Treatment is binary, and either on or off, covering the entire domain. For the purposes of this model, we are only considering treatment to be cytostatic, preventing cells from dividing, representing a cytostatic therapy analogous to tyrosine kinase inhibitors (TKIs) used in non-small cell lung carcinoma. The carrying capacity for the ODE is 10000 cells, and the domain size of the ABM 2D grid is 10000 spaces. In the ABM, the initial population is randomly placed on the grid. Each agent in the ABM followed the simple rule set seen in **ABM Decision Tree**. Additionally, the agents in the ABM utilized a Moore neighborhood for decision making.

Each mutation incrementally increases fitness relative to the fully resistant phenotype. The number of mutational steps between the initial persister state and full resistance is varied across simulations (1, 2, 5, 10, 15, or 20 steps), enabling exploration of evolutionary trajectories across different adaptive landscapes.

**Agent-Based Model Implementation**

The ABM is defined on a two-dimensional grid of 10,000 lattice sites, each representing an individual spatial location that can be occupied by a single cell. Agents interact with their local environment through a Moore neighborhood, consisting of the eight adjacent lattice sites. The initial population is randomly seeded across the domain, with relative proportions of sensitive, persister, and mutator cells determined by experimental or theoretical assumptions (Supplementary Table 1). At each discrete time step, agents update asynchronously according to probabilistic rules governing division, death, and mutation (Fig. S1). The ABM was executed for 50 independent stochastic realizations to account for random variation and to quantify variability across simulation runs.

**Ordinary Differential Equation (ODE) Model**

To complement the ABM and isolate the effects of stochasticity and spatial structure, a corresponding deterministic ordinary differential equation (ODE) model was constructed under identical biological assumptions and rate parameters. The ODE system models population-level dynamics without spatial constraints and assumes a total carrying capacity of 1,000 cells. Unlike the ABM, which captures local competition and stochastic evolutionary paths, the ODE model produces a smooth deterministic trajectory for comparison.

**Model parameters:**

The model parameters were informed from experimental data and from a literature review. Identical parameters and rates were shared between the ABM and the ODE, when possible.

To model cytostatic conditions (Fig. S2A), the proliferation term $G_s(1 - N/K)$ was reduced while the death rate *D* remained constant. To model cytotoxic conditions (Fig. S2B) the death rate *D* was increased while the proliferation term $G_s(1 - N/K)$ remained constant. All other parameters were unchanged from the base model used in Fig. 2C.

**Model implementation:**

Both models were designed and implemented in the Python programming language. Python was also used for all data processing and visualization.

**Model validation:**

To validate the models, we compared their predictions to empirical data and used sensitivity analysis to explore the robustness of the results. The models were run for a set number of iterations, and the results were compared between both models. These results were then used to inform and predict in vitro experiments.

**ABM and ODE Parameters**

| Parameter | Value | Source |
|---|---|---|
| Sensitive population growth rate | 0-1 | Experimental Data |
| Intermediate population growth rate | 0-1 | Experimental Data |
| Fitness of the resistant population | 1 | Experimental Data |
| Death rate | 0.25 | Experimental Data |
| Mutation rate | 0.009 | Experimental Data |
| Number of steps towards resistance | 1 - 20 | Experimental Data |

| | Parameters |
|---|---|
| $t$ | Time (days) |
| $K$ | Carrying capacity |
| $N$ | Total Population |
| $G_S$ | Sensitive population growth rate |
| $g_n$ | Intermediate population growth rate |
| $f_R$ | Fitness of the resistant population |
| $D$ | Death rate |
| $\mu$ | Mutation rate |
| $S$ | Sensitive population |
| $m$ | Mutator population |
| $p$ | Non-mutator population |
| $R$ | Resistant population |
| $A$ | Total number of steps towards resistance |
| $n$ | Number of intermediate steps towards resistance |

### ABM Decision Tree

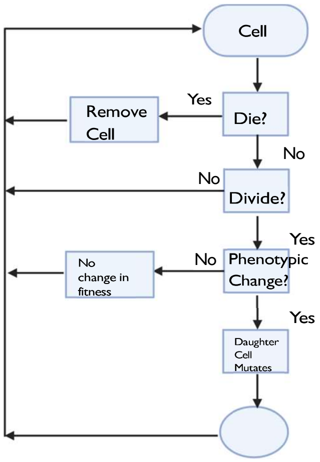


### ODE System of Equations

$$\frac{dS(t)}{dt} = S\left(t\right) \cdot \left(G_S\left(1 - \frac{N}{K}\right) - D\right)$$

$$\frac{dm_n(t)}{dt} = \frac{dp_n(t)}{dt} = U_n \cdot \left(g_n\left(1 - \frac{N}{K}\right) - D - \mu_U\right) + \left(U_{n-1} \cdot \mu_U\right),$$

$$g_n = \frac{n}{A} \cdot f_R + \frac{(A-n)}{A} \cdot D$$

$$\frac{dR_m}{dt} = \frac{R_p}{dt} = R_U \cdot \left(f_R\left(1 - \frac{N}{K}\right) - D\right) + \left(U_{A-1} \cdot \mu_U\right)$$
